# Functional connectomics spanning multiple areas of mouse visual cortex

**DOI:** 10.1101/2021.07.28.454025

**Authors:** The MICrONS Consortium, J. Alexander Bae, Mahaly Baptiste, Caitlyn A. Bishop, Agnes L. Bodor, Derrick Brittain, JoAnn Buchanan, Daniel J. Bumbarger, Manuel A. Castro, Brendan Celii, Erick Cobos, Forrest Collman, Nuno Maçarico da Costa, Sven Dorkenwald, Leila Elabbady, Paul G. Fahey, Tim Fliss, Emmanouil Froudarakis, Jay Gager, Clare Gamlin, William Gray-Roncal, Akhilesh Halageri, James Hebditch, Zhen Jia, Emily Joyce, Justin Joyce, Chris Jordan, Daniel Kapner, Nico Kemnitz, Sam Kinn, Lindsey M. Kitchell, Selden Koolman, Kai Kuehner, Kisuk Lee, Kai Li, Ran Lu, Thomas Macrina, Gayathri Mahalingam, Jordan Matelsky, Sarah McReynolds, Elanine Miranda, Eric Mitchell, Shanka Subhra Mondal, Merlin Moore, Shang Mu, Taliah Muhammad, Barak Nehoran, Oluwaseun Ogedengbe, Christos Papadopoulos, Stelios Papadopoulos, Saumil Patel, Xaq Pitkow, Sergiy Popovych, Anthony Ramos, R. Clay Reid, Jacob Reimer, Patricia K. Rivlin, Victoria Rose, Casey M. Schneider-Mizell, H. Sebastian Seung, Ben Silverman, William Silversmith, Amy Sterling, Fabian H. Sinz, Cameron L. Smith, Shelby Suckow, Marc Takeno, Zheng H. Tan, Andreas S. Tolias, Russel Torres, Nicholas L. Turner, Edgar Y. Walker, Tianyu Wang, Adrian Wanner, Brock A. Wester, Grace Williams, Sarah Williams, Kyle Willie, Ryan Willie, William Wong, Jingpeng Wu, Chris Xu, Runzhe Yang, Dimitri Yatsenko, Fei Ye, Wenjing Yin, Rob Young, Szi-chieh Yu, Daniel Xenes, Chi Zhang

## Abstract

To understand the brain we must relate neurons’ functional responses to the circuit architecture that shapes them. Here, we present a large functional connectomics dataset with dense calcium imaging of a millimeter scale volume. We recorded activity from approximately 75,000 neurons in primary visual cortex (VISp) and three higher visual areas (VISrl, VISal and VISlm) in an awake mouse viewing natural movies and synthetic stimuli. The functional data were co-registered with a volumetric electron microscopy (EM) reconstruction containing more than 200,000 cells and 0.5 billion synapses. Subsequent proofreading of a subset of neurons in this volume yielded reconstructions that include complete dendritic trees as well the local and inter-areal axonal projections that map up to thousands of cell-to-cell connections per neuron. Here, we release this dataset as an open-access resource to the scientific community including a set of tools that facilitate data retrieval and downstream analysis. In accompanying papers we describe our findings using the dataset to provide a comprehensive structural characterization of cortical cell types^1–3^ and the most detailed synaptic level connectivity diagram of a cortical column to date^2^, uncovering unique cell-type specific inhibitory motifs that can be linked to gene expression data^4^. Functionally, we identify new computational principles of how information is integrated across visual space^5^, characterize novel types of neuronal invariances^6^ and bring structure and function together to decipher a general principle that wires excitatory neurons within and across areas^7, 8^.

## Introduction

Francis Crick wrote in 1979^9^ that “*It is no use asking for the impossible, such as, say, the exact wiring diagram for a cubic millimeter of brain tissue and the way all its neurons are firing*.” Crick’s impossible request was motivated by the idea that the function of every neuron depends strongly on the functions of its synaptically connected partners, as already hypothesized by the earliest studies of visual cortex^10^. Neuroscientists have long been interested in this dependence but while the physiology of single neurons has been linked to morphological and connectivity properties^11–14^, relating functional properties of multiple neurons to the exact connectivity between them has been far more difficult. This pursuit of inferring both function and connectivity of a network was initially attempted using electrophysiology recordings^15, 16^ or optical imaging combined with anatomy^17–19^

More recently, multi-photon calcium imaging has allowed ever larger and more densely sampled groups of neurons to be studied *in vivo*^20–23^. These recordings have been correlated with connectivity, using *in vitro* electrophysiological^24, 25^ and viral methods ^26, 27^. Much has been learned from these kinds of experiments, but they only provide fragmentary information about neural activity and connectivity, rather than the comprehensive information that Crick dreamed of. Steps towards this goal have been achieved with volumetric electron microscopy combined with calcium imaging, an approach known as functional connectomics, in the visual cortex^28–31^, mammalian retina^32–34^, and several other systems ^35, 36^.

Here we report the results of a five-year multi-institutional collaboration where we applied calcium imaging *in vivo* to record the responses to visual stimuli of a large fraction of the pyramidal neurons in a cubic millimeter of mouse visual cortex (1.3×0.87×0.82 mm^3^ *in vivo* dimensions). The same cubic millimeter was then imaged with serial section electron microscopy (EM), generating over a petabyte of image data. The EM images were processed by convolutional nets to automatically reconstruct neurons and synapses in 3D. Finally, the 3D reconstruction was co-registered with the calcium images, enabling the visual responses of neurons to be matched with their connectivity. We have released the data as a public resource (www.microns-explorer.org). The resource contains the visual responses of up to 75,909 cells, one of the largest collections of neural recordings obtained from a single animal. The resource also includes a neuronal wiring diagram, which is becoming increasingly complete and accurate as proofreading of the automated reconstruction proceeds.

This work represents the largest functional connectomics dataset by volume thickness, number of cells, and number of synapses detected. It contains tens of thousands of complete dendritic arbors and the longest mammalian axonal arbors reconstructed by EM. The dataset also contains multiple cortical visual areas enabling both intra-and inter-areal connectivity analysis. We will refer to these primary data and derived data products as the *MICrONS multi-area dataset*, as they are the results of an IARPA program named “Machine Intelligence from Cortical Networks.” Several of the technologies used, including imaging, proofreading, segmentation, and analysis infrastructure, were initially piloted on a smaller dataset^30, 31, 37^ before being scaled up for the present work.

This public resource (Fig 1) includes pyramidal neurons from all layers (underlined text contains links to public data), such as <u>L5 thick tufted</u>, <u>L5 NP</u>, <u>L4</u>, and <u>L2/3</u>. It includes inhibitory neurons from many classes, such as <u>bipolar cells</u>, <u>basket cells</u>, a <u>chandelier cell</u>, and <u>Martinotti cells</u>. It also includes non-neuronal cells, such as <u>astrocytes</u> and <u>microglia</u>. The network of <u>blood</u> <u>vessels</u> is segmented, and contains some erroneously merged astrocytes. Using the interactive visualization tools, one can visualize the <u>input and output synapses of a single cell</u>. T<u>he</u> <u>database of functional recordings</u> is also available for download to explore how cells responded to visual stimuli. For a subset of the neurons, we have generated a correspondence between their functional recording and their anatomical reconstruction.

**Figure 1.**
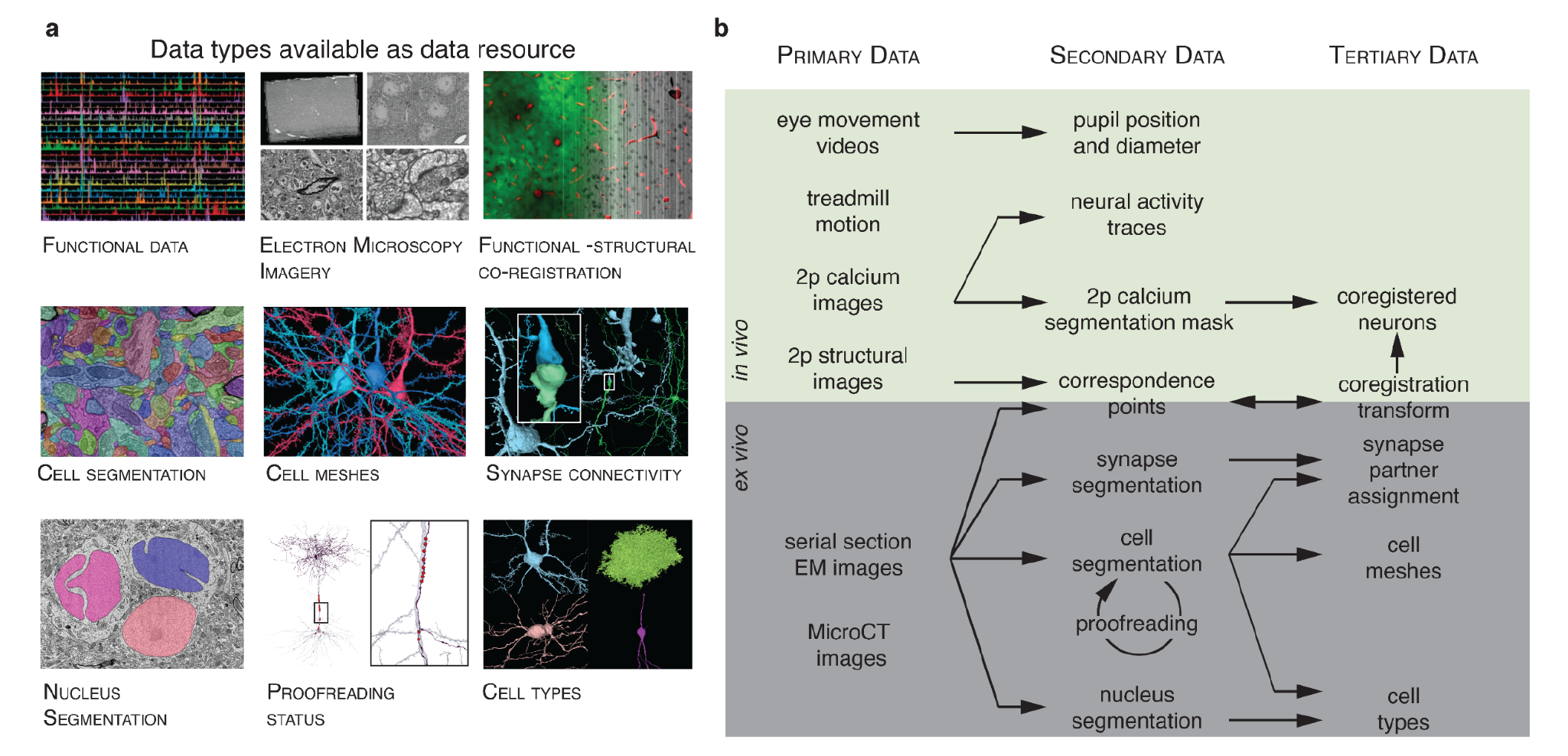
Resource Data Type and Data products. **a)** The nine data resources publicly available in www.microns-explorer.com. **b)** Relationship between different data types. The primary *in vivo* data resource consists of two-photon calcium images, two-photon structural images, natural videos and parametric stimuli used as visual input, and behavioral measurements. The secondary (i.e. derived) *in vivo* data resource includes the responses of approximately 75,909 pyramidal cells from cortical layer 2 (L2) to 5 (L5) segmented from the calcium movies, along with the pupil position and diameter extracted from the video of eye movements and locomotion measured on a single-axis treadmill. The primary anatomical data are composed of *ex vivo* serial section transmission EM images registered with the *in vivo* two photon structural stack. The volume includes a portion of primary visual cortex (VISp) and higher visual areas VISlm, VISrl, and VISal ^52^ for all cortical layers except extremes of L1. The secondary anatomical data is derived from the serial section EM image stack, and consists of (a) semi-automated segmentation of cells, (b) automated segmentation of nuclei, and (c) automatically detected synapses. The tertiary anatomical data consists of (a) assignments of the synapses to presynaptic and postsynaptic cells, (b) triangle meshes for these segments, (c) classification of nuclei as neuronal versus non-neuronal, and (d) classification of neurons into excitatory and inhibitory cell classes. Secondary data for coregistration of *in vivo* and *ex vivo* images consists of manually chosen correspondence points between two-photon structural images and EM images. Tertiary coregistration data is a transformation derived from these correspondence points. The transformation is then used to facilitate the matching of cell indices between the two-photon calcium cell segmentation masks and the EM segmentation cells

This manuscript presents a comprehensive description of the data, along with the tools to explore it, in order for others to make use of it in neuroscience or in other fields ^38^. The first scientific findings to emerge from the data are described in the accompanying manuscripts ^1–4, 7^. Here we confine ourselves to general remarks about the scientific implications. Some manuscripts characterize cortical cell types and their connectivity ^1–3^, applying EM data to renew a tradition dating back to Cajal. This is timely because a major endeavor in contemporary neuroscience is to systematically map the properties of brain cells. One prominent technology has been single cell RNA sequencing, which is being used to create taxonomies of brain cell types defined by transcriptomes ^39, 40^. Therefore accompanying manuscripts also begin to establish correspondences between the connectivity of anatomically defined cell types with transcriptomically defined types^4^. A major product of our work is the rules by which cell types are wired to each other to form neuronal circuits^2, 4^.

In this respect, our work parallels another milestone of connectomics, the imminent completion of the *Drosophila* connectome ^41–43;^ only 20% of the neuron types described in the EM connectome of the central brain were previously described in the literature ^41^. There is however an important difference to be drawn with *Drosophila*, in which a cell type often consists of just a few neurons that share similar functional properties that are reproducible across individuals. Owing to this stereotypy, a connectome mapped in one fly can usually be used by researchers studying neuronal function in other flies. Rules of connectivity based on cell types have proved sufficient for understanding and modeling many functions of increasingly complex neural circuits ^44–46^. Conversely, a single cell type in a mammalian brain encompasses a huge number of cells, which generally exhibit different tuning preferences. This is why it is important to combine cortical connectomics with functional studies of the same neurons in the same brain. And this is why the mapping of cortical connectivity must go beyond rules that depend solely on cell types.

Furthermore, advances in large scale recording technologies have made it possible to explore at scale functional response properties of mammalian neurons beyond simple parametric tuning curves ^22, 47–49^. Comprehensively capturing this broader diversity of neuronal responses necessitates reconsidering the models and paradigms we commonly use to characterize them. As one powerful approach, an accompanying paper builds a functional digital twin of the MICrONS multi-area dataset using deep learning methods leveraging a “foundation model” of the mouse visual system which was trained using large scale data sets from multiple visual cortical areas and mice ^7, 8^. This model generalizes well, accurately predicting neuronal responses not only to natural videos, but also to various new stimulus domains, such as coherent moving dots and noise patterns, as confirmed through in vivo testing^7, 8^. This model serves as a valuable tool to comprehensively analyze the relationships between circuit function and structure that can help build a mechanistic understanding of cortical computations ^5–7^.

We also note that our cubic millimeter volume, while tiny compared to an entire mouse brain, is over an order of magnitude larger than a *Drosophila* brain. Some of the accompanying manuscripts highlight the tools and techniques which were invented to scale up connectomics to a cubic millimeter ^8, 50, 51^. These technologies are already having a broader impact by enabling connectomics for many brain regions and multiple species. For example, the same image processing pipeline was already repurposed for the forthcoming *Drosophila* connectome, the first adult connectome to be completed since that of *C. elegans*. Further development of the MICrONS technologies should eventually render Crick’s dream not only possible, but routine.

## Results

### Overview

The data were collected from a single animal and involved a pipeline spanning three primary sites. First, two-photon *in vivo* calcium imaging under various visual stimulation conditions was performed at Baylor College of Medicine. Then the animal was shipped to the Allen Institute, where the imaged tissue volume was extracted, prepared for EM imaging, sectioned, and imaged over a period of 6 months of continuous imaging. The EM data were then montaged, roughly aligned, and delivered to Princeton University, where fine alignment was performed and the volume was densely segmented. Finally, extensive proofreading was performed on a subset of neurons to correct errors of automated segmentation, and cell types and various other structural features were annotated (Fig 2).

**Figure 2.**
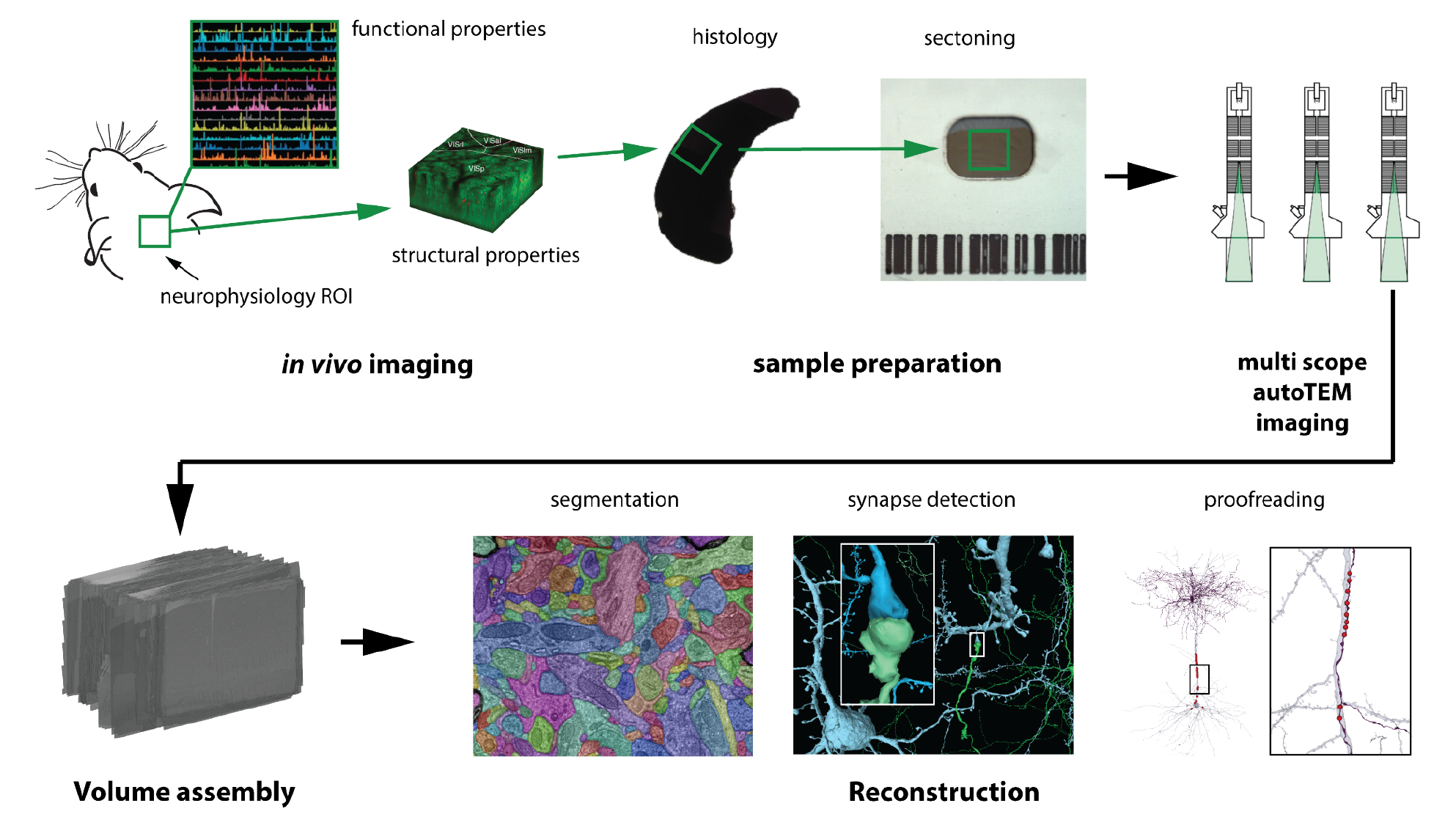
Major experimental steps in the data acquisition workflow.

### Functional Data

#### Two-Photon Calcium Imaging

The calcium-imaging data includes the responses to visual stimuli of an estimated 75,909 excitatory neurons spanning cortical layers 2 through 5 across four visual areas, in a transgenic mouse expressing GCaMP6s in excitatory neurons via SLC17a7-Cre^53^ and Ai162^54^. The dataset contains fourteen individual scans, collected between P75 and P81, spanning a volume of approximately 1200 *x* 1100 *x* 500 µm^3^ (anteroposterior *x* mediolateral *x* radial depth, Fig 3a). The center of the volume was placed at the junction of primary visual cortex (VISp) and three higher visual areas, lateromedial area (VISlm), rostrolateral area (VISrl) and anterolateral area (VISal) in order to image retinotopically-matched neurons connected via inter-areal feedforward and feedback connections.

**Figure 3.**
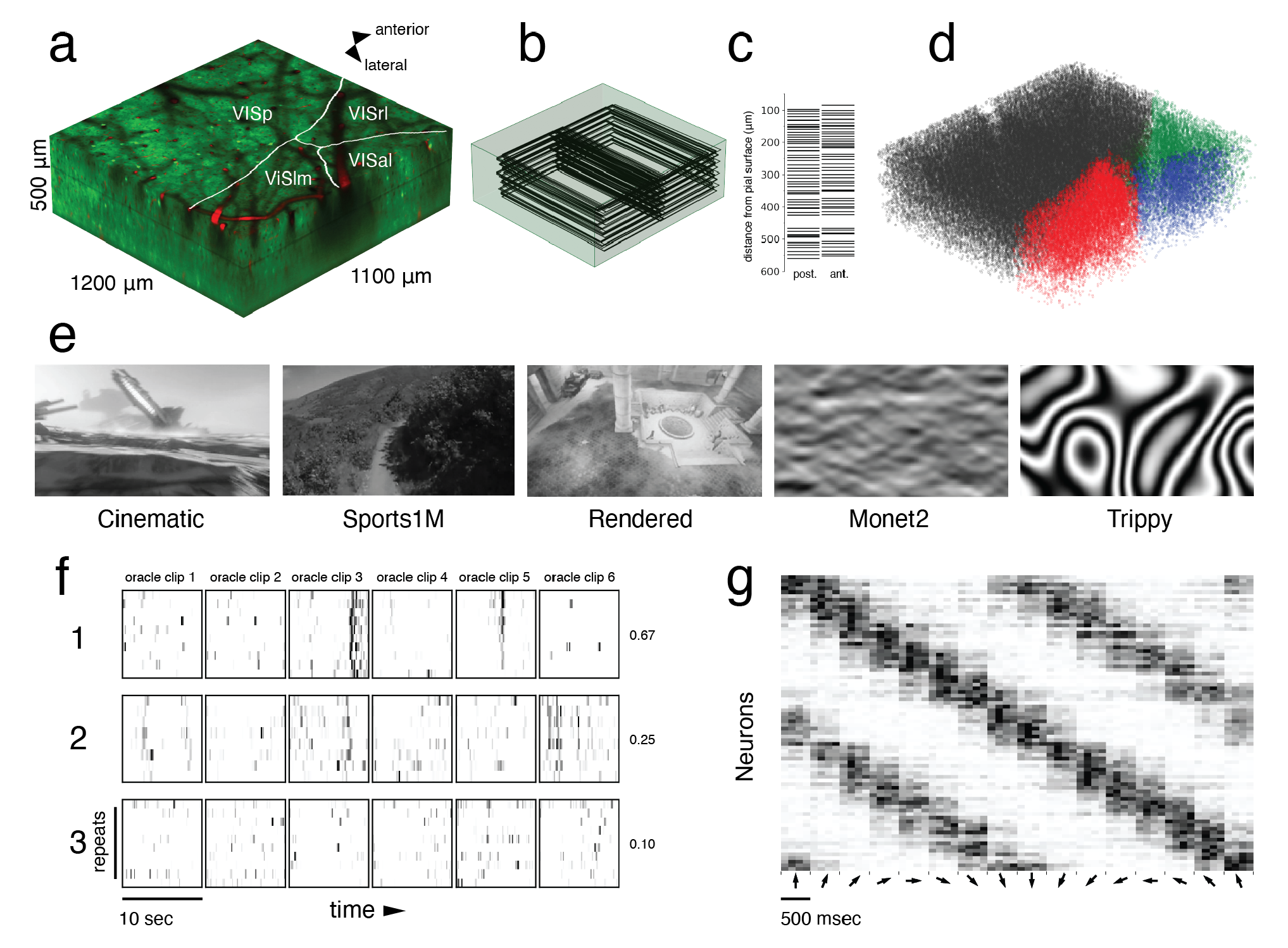
*In Vivo* Calcium Imaging Data. **a)** Representation of the two-photon functionally imaged volume with area boundaries (white) and vascular label from structural stack (red). **b)** Wireframe representation of 104 planes registered in the structural two-photon stack. **c)** Mean depth of each registered field relative to the pial surface. Left: posterior fields, Right: anterior fields **d)** 3D scatter plot of each functional mask in its registered location in the structural two-photon stack (VISp: black, VISlm: red, VISal: blue, VISrl: green). **e)** Example frames from each of the five stimulus types (Cinematic, Sports1M, Rendered, Monet2, Trippy) shown to the mouse. **f)** Raster of deconvolved calcium activity for three neurons to repeated stimulus trials (oracle trials; 6 sequential clips x 10 repeats, with each repeat normalized independently). (1) high, (2) medium, and (3) low oracle scores (right). **g)** Trial averaged raster of deconvolved calcium activity for 80 neurons to 40 Monet2 trial (16 randomly ordered directions) grouped by preferred direction (5 neurons/ direction; alternating blue shading) and sorted according to the stimulus directions. Central 500 msec of trial average raster for each direction (out of 937 msec)

Each scan consisted of two adjacent overlapping 620 µm-wide fields at multiple imaging planes, imaged with the wide field-of-view of the two-photon random access mesoscope (2P-RAM)^55^. The scans ranged up to approximately 500 µm in depth, with a target spacing of 10-15 µm to maximize the coverage of imaged cells in the volume (Fig 3b,c). For eleven of the fourteen scans, four imaging planes were distributed widely in depth using the mesoscope remote focus, spanning roughly 300-400 µm with an average spacing of approximately 125 µm between planes for near-simultaneous recording across multiple cortical layers. In the remaining three scans, fewer planes were imaged at 10-20 µm spacing to achieve a higher effective pixel density. These higher resolution scans were designed to be amenable to future efforts to extract signals from large apical dendrites from deeper layer 5 and layer 6 neurons, for example using EM-Assisted Source Extraction (EASE^56^). However, for this release, imaging data was automatically segmented only from somas using a constrained non-negative matrix factorization approach and fluorescence traces were extracted and deconvolved to yield activity traces ^57^. In total, 125,413 masks were generated across fourteen scans, of which 115,372 were automatically classified as somatic masks by a trained classifier (Fig 3d^57^).

The functional data collection relied on newly-established technologies, especially the 2P-RAM mesoscope developed at Janelia^55^. In addition, we developed an imaging workflow with the goal of full coverage within the target volume. This required several optimizations, for example in order to densely target scan planes across multiple days, we needed a common reference frame to assess the coverage of scans within the volume. Therefore, in addition to the functional scans, high-resolution (0.5 - 1 px/µm) structural volumes were acquired for registration with the subsequent EM data. At the end of each imaging day, individual imaging fields of the functional scans were independently registered into a structural stack (Fig 3b,c). This allowed us to target scans in subsequent sessions to optimize coverage across depth. On the last day of imaging, a two-channel (green, red) structural stack was collected at 0.5 px/um after injection of fluorescent dye (Texas Red) to label vasculature, enhancing fiducial labeling for co-registration with the EM volume (Fig 3a).

After registration of the functional imaging field with the structural stack, 2D centroids from the segmentation were assigned 3D centroids in the shared structural stack coordinate space, based on a greedy assignment of 3D proximity. Based on this analysis, we estimate the functional imaging volume contains 75,909 unique functionally-imaged neurons consolidated from 115,372 segmented somatic masks, with many neurons imaged in two or more scans.

#### Behavioral Monitoring and Visual Stimulation

During imaging, the animal was head-restrained, and the stimulus was presented to the left visual field. Treadmill rotation (single axis) and video of the animal’s left eye were captured throughout the scan, yielding the locomotion velocity, eye movements, and pupil diameter data included here.

The stimulus for each scan was approximately 84 minutes, and consisted of natural and parametric movie stimuli. The majority of the stimulus (64 min) was made up of 10 second clips drawn from cinema, the Sports-1M^58^ dataset, or rendered first-person (Point of View) POV movement through a virtual environment (Fig 3e). Our goal was to approximate natural statistical complexity to cover a sufficiently large feature space. These data can support multiple lines of investigation including applying deep-learning-based systems identification methods to build highly accurate models that predict neural responses to arbitrary visual stimuli ^59^ ^22^ ^60^. These models enable a systematic characterization of tuning functions with minimal assumptions relative to classical methods using parametric stimuli^22^.

The stimulus composition included a mixture of unique stimuli for each scan, some that were repeated across every scan, and some that were repeated within each scan. In particular, six natural movie stimuli clips totalling one minute were repeated in the same order 10 times per scan, and were used to evaluate the reliability of the neural responses to repeated visual stimuli (Fig 3f). This "oracle score" serves as an important quality metric since reliable responses are not observed when imaging conditions are poor or the animal is not engaged with the stimulus

To relate our findings to previous work, we also included a battery of parametric stimuli ("Monet2" and "Trippy", 10 minutes each) that were generated to produce spatially decorrelated stimuli suitable for characterizing receptive fields, while also containing local or global directional and orientation components for extracting basic tuning properties such as orientation selectivity (Fig 3e,g).

### The Electron Microscopy volume

After the *in vivo* neurophysiology data collection, we imaged the same volume of cortex *ex vivo* using Transmission Electron Microscopy (TEM) which allowed us to map the connectivity of neurons for which we measured functional properties. These required considerable scaling from previous state of the art datasets, with particular emphasis on automation^61^ and on reducing rare but potentially catastrophic events that could incur multiple serial section loss.

The tissue sample was trimmed and sectioned into 27,972 serial sections (nominal thickness: 40 nm) onto grid tape to facilitate automated imaging ^61, 62^. Although the cutting was automated, it was supervised by humans who worked in shifts around the clock for 12 days. They were ready to stop and restart the ultramicrotome at a moment’s notice if there was a risk of multiple section loss. As will be described later, the EM dataset is subdivided into two subvolumes due to sectioning and imaging events (see methods for details of sectioning timeline and artifacts). A total of 26,652 sections were imaged by a fleet of five customized automated Transmission Electron Microscopes (autoTEMs^61^), which took approximately 6 months to complete and produced a dataset composed of 2 PB of raw data at a ∼4nm resolution (Fig 4d-h).

**Figure 4.**
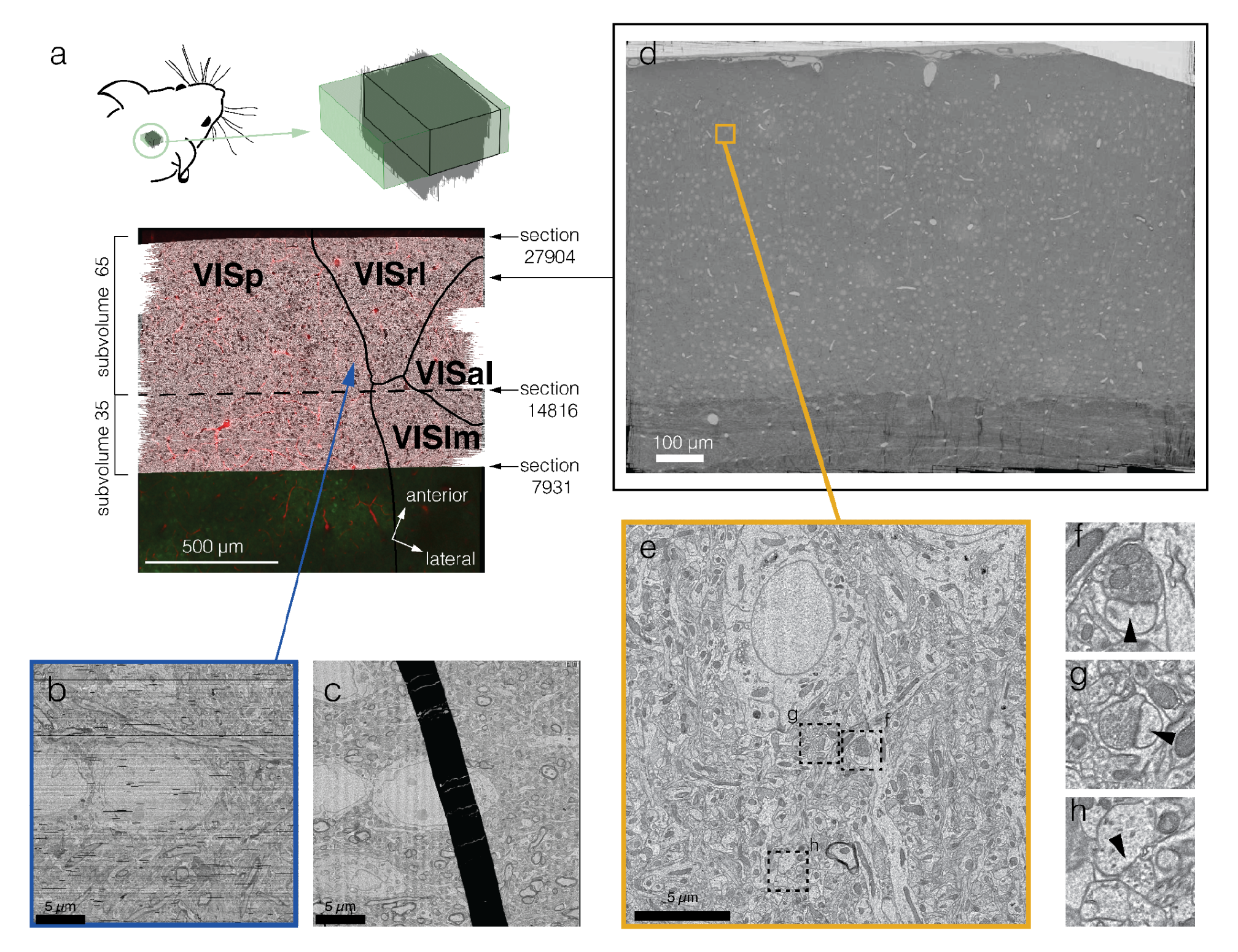
EM Dataset. **a)** Top view of EM dataset (gray) registered with the *in vivo* 2-photon structural dataset (vasculature in red and GCaMP in green). Area borders calculated from calcium imaging are shown as black lines (VISp, primary visual cortex; VISrl, rostro lateral visual cortex; VISal anterolateral visual cortex; VISlm lateromedial visual cortex). The two portions of the dataset are separated by a dashed line. **b)** Top view of small region showing the quality of the fine alignment and its robustness to large folds shown in “**c”** (link to dataset: https://ngl.microns-explorer.org/#!gs://microns-static-links/mm3/data_fig/4b.json). **d)** Montage of a single section showing the coverage from pia to white matter and across 3 different cortical regions. **e)** Example of a single tile from the section in ”**d”** with dashed squares representing the location in **f - h**. **f-g)** examples of excitatory synapses indicated with arrowheads (link to f in dataset: https://ngl.microns-explorer.org/#!gs://microns-static-links/mm3/data_fig/4f.json; link to g in dataset:https://ngl.microns-explorer.org/#!gs://microns-static-links/mm3/data_fig/4g.json) **h)** examples of inhibitory synapse indicated with arrowhead (link to h in dataset: https://ngl.microns-explorer.org/%23%21gs://microns-static-links/mm3/data_fig/4h.json).

An 800 µm region (sections 7,931–27,904) (Fig 4a) was selected for further processing, as it had no consecutive section loss and an overall section loss of around 0.1%. This region contains approximately 95 million individual tiles that were stitched into 2D montages per section and then aligned in 3D. Due to the re-trimming of the block and the requirement for a knife change (see methods), the EM data are divided into two subvolumes (Fig 4a). One subvolume contains approximately 35% of the sections (sections 7,931–14,815) and the other 65% containing sections 14,816–27,904. The two subvolumes were processed individually and later fine aligned to each other in the same global coordinate frame, allowing the tracing of axons and dendrites across their border (Fig 5). To facilitate the reconstruction process across the division between the two subvolumes, a composite image of the partial sections was created at the interface. However, the two subvolumes were reconstructed separately and each has a distinct representation in the analysis infrastructure and database.

**Figure 5.**
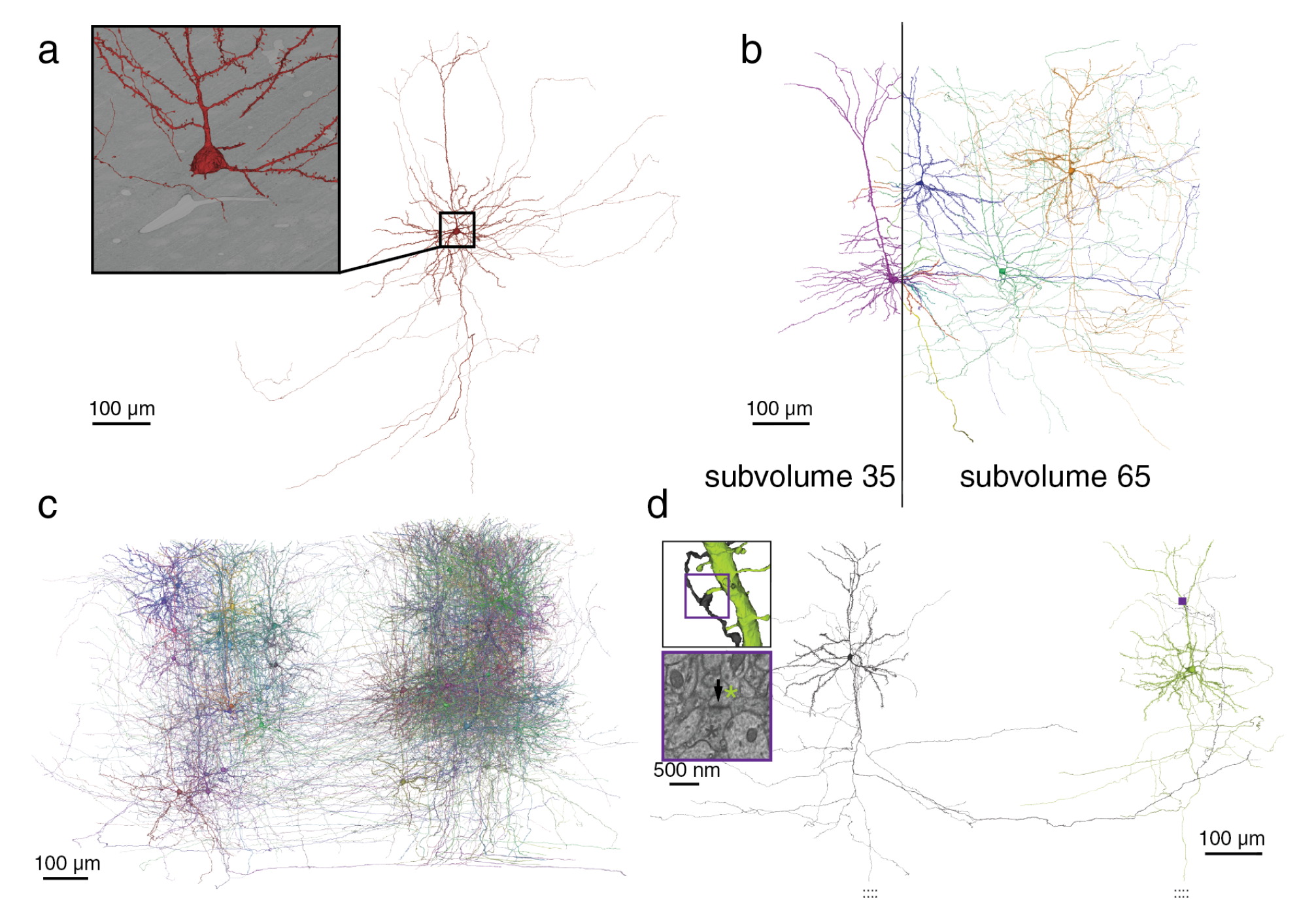
Reconstruction. **a)** A pyramidal cell as it is reconstructed from the EM imagery (inset). **b)** Pyramidal cells from both subvolumes as they cross the subvolume boundary. **c)** All 78 proofread pyramidal cells from subvolume 65. **d)** A distant pair of pyramidal cells connected by a synapse within subvolume 65.

Accurate reconstruction requires extremely accurate stitching and alignment of images hundreds of thousands of pixels on a side. To achieve this at petabyte scale, we split the process into distinct coarse and fine pipelines. For the coarse pipeline, sections were initially stitched using a per-image affine transformation, and a polynomial transformation model was applied to a subset of sections whose stitching quality had a local misalignment error above5 pixels^63^. Downsampled 2D stitched sections were then roughly aligned in 3D. The rough alignment process ensured global consistency within the dataset and accounted for images from multiple autoTEMs with varied image sizes and resolutions. It also corrected for locally varying misalignments such as scale differences and deformations between sections and aids the fine alignment process.

To further refine image alignment, we developed a set of convolutional networks to estimate pixelwise displacement fields between pairs of neighboring sections^51^. This process was able to correct nonlinear misalignments around cracks and folds that occurred during sectioning. Though this fine alignment does not restore the missing data inside a fold, it was still effective in correcting the distortions caused by large folds (Fig 4b-c), which caused large displacements between sections and were the main cause of reconstruction errors. While imaging was performed with 4 nm resolution, the aligned imagery volume was generated at 8 nm resolution to address file size considerations

### Automated reconstruction of cellular processes

We densely segmented cellular processes across the volume using affinity-predicting convolutional neural networks and mean-affinity agglomeration (Fig 5a)^31, 64–66^. Segmentation was not attempted where the alignment accuracy was deemed insufficient or tissue was missing or occluded over multiple sections.

The automatic segmentation produced highly accurate dendritic arbors before proofreading, allowing for morphological identification of broad cell types. Most dendritic spines are properly associated with their dendritic trunk. Recovery of larger caliber axons, those of inhibitory neurons, and the initial portions of excitatory neurons was also typically successful. Due to the high frequency of imaging defects in the shallower and deeper portions of the dataset, processes near the pia and white matter often contain errors. Many non-neuronal objects are also well-segmented, including astrocytes, microglia, and blood vessels. The two subvolumes of the dataset were segmented separately, but the alignment between the two is sufficient for manually tracing between them (Fig 5b).

Nuclei were also automatically segmented (n=144,120) within subvolume 65 using a distinct convolutional network^51^. To use nucleus shape to map cell classes across the dataset, we manually labeled a subset of the 2,751 nuclei in a 100 µm square column of the dataset as non-neuronal, excitatory or inhibitory. We then developed machine learning models to automate the detection of neurons, as well as classify cells at different levels of resolution ^1, 50^ within the subvolume with high accuracy (see Methods). The results of this nucleus segmentation, manual cell classification and model building are as part of this data resource.

Synaptic contacts were automatically segmented in the aligned EM image, and the presynaptic and postsynaptic partners from the cell segmentation were automatically assigned to identify each synapse (Fig 5d)^31, 64–66^. We automatically detected and associated a total of 524 million synaptic clefts across both subvolumes (subvolume 35: 186 million, subvolume 65: 337 million). We manually identified synapses in 70 small subvolumes (n=8,611 synapses) distributed across the dataset, giving the automated detection an estimated precision of 96% and recall of 89% (Supplemental Fig 1) ^67^ with a partner assignment accuracy of 98%.

### Proofreading

While the automated segmentation creates impressive reconstructions, proofreading is required to make those reconstructions more complete and accurate. The proofreading process involves both merging additional segments of the neurons that were missing in the reconstruction, and splitting segments that were incorrectly associated with a neuron. To do real-time collaborative proofreading in a petascale dataset, we employed the ChunkedGraph proofreading system^31^ that can be used with a modified version of Neuroglancer as a user interface or a REST API for computationally-driven edits. This flexibility enabled the proofreading methods to be tailored to different scientific needs, including manual, semi-automated, and automated proofreading. Note that all proofreading has happened in subvolume 65.

The segmentation released here contains all edits of the proofreading that had occurred as of April 6, 2023. Proofreading was performed by individual scientists and focused teams of proofreaders to both support targeted scientific discovery for companion studies ^1–4, 7^ and to correct errors that most impacted general connectivity. Because of this, the level of completeness differs across these cells (Fig 5), as neurons have been proofread as part of multiple MICrONS data analysis projects^1–4, 7^. For example, in the functional connectomics study, we proofread the full extent of axonal and dendritic arbors of 78 excitatory neurons within subvolume 65 (Fig 5c), while for a broad columnar sample only the dendrites of 1,188 excitatory neurons were proofread. The result is a wide variation in edits per neuron with more edits generally corresponding to more extensive axons (Supplemental Fig 2). The released dataset includes 601 neurons that have been extensively proofread, with 46,241 splits and 38,694 merges to eliminate any incorrect merges and extend many false splits. As a result, for these particular neurons all synapses - both input and output - are now correctly associated. The most time consuming task is extending axons, and thus this is where the data varies most across cells and studies. The fully reconstructed excitatory neurons mentioned above contain some of the most extensive axonal arbors reconstructed in the neocortex at EM resolution (see discussion), with excitatory axon lengths ranging from 2.5 to 18.9 mm and a mean of 714 synaptic outputs (range: 192-1,893) and inhibitory axons ranging from 1.1 to 32.3 mm and a mean of 2,828 synaptic outputs (range: 75–13,956). For most other companion studies, axonal arbors were typically partially extended (e.g. extensively but not completely extending inhibitory axons) or limited by the time spent on each incomplete segment end. In general, inhibitory axons were more complete in the automated reconstruction, likely due to their axons being slightly thicker than most excitatory axons.

In addition to proofreading axons and dendrites, we made widespread edits to enhance the general dataset quality. Following the automated segmentation, there were 7,050 segmented objects consisting of a total of 17,753 neurons that were merged together (based on nucleus segmentation), preventing analysis of these cells. Using a combination of manual and automated error detection workflows, we have split 99.4% (n=7,011) of such merged objects, bringing the total number of individually segmented neurons to 81,961 (Supplemental Fig 3). To work through such dataset-wide tasks more quickly, we developed and validated an automated error detection and correction workflow using graph and morphological analysis to not only identify merge error locations, but also generate edits that could be executed using PyChunkedGraph (PCG)^31^. This automated approach (NEURD) was also used to remove false axon merges onto dendritic segments and split axon branches with abnormally high degree across the dataset, totalling more than 164,000 edits^50^.

### Functional - Structural Co-registration

Functional connectomics requires that cells are matched between the two-photon calcium imaging and EM coordinate frames. This was achieved by a three-phase approach using expert annotations and automatic methods. In the first step we generated a co-registration transform using a set of 2,934 expert matched fiducials between the EM volume and the two-photon structural dataset (1994 somata and 942 blood vessels, mostly branch points, which are available as part of the resource, see Methods). To evaluate the error of the transform we evaluated the distance in microns between the location of a fiducial after co-registration and its original location; a perfect co-registration would have residuals of zero microns). The average residual was 3.8 microns.

For the second step we used the results of the transform to guide a group of experts to manually validate matches from 13,925 functional ROIs to 12,054 individual EM neurons to yield high-confidence matches for analysis and provide "ground truth" for a fully-automated approach. These results help validate the first phase as most matched ROIs have low residuals and high separation scores (Supplemental Fig 4). Furthermore, ROIs with at least moderate visual responses that are matched to the same neuron have higher signal correlations than nearby neurons (Supplemental Fig 4). In the third and final step, we used fully automated methods to match an additional 62,378 functional ROIs with residuals below 15µm matched to 34,063 EM neurons, shared publicly as apl_functional_coreg_forward_v5. Briefly, a segmentation of the blood vessels was used to refine the initial co-registration with a fine-scale deformable registration between the EM and the two photon in vivo imaging space (see methods). This automated method produced results for 99.5% of the manually matched ROIs. Considering all of the matches, the automated method achieved 73.8% agreement with the manual matchers, but application of a restriction that only considered matches with separation above 6µm and residual below 15µm increased the level of agreement to 91.4%, while including 20,329 functional ROIs and 14,829 EM neurons. (Supplemental Fig 5).

### Integrated Analysis

In order to create a resource for the neuroscience community, we have made the data from each of the steps described above — functional imaging, the EM subvolumes, segmentation, and a variety of annotations — publicly available on the MICrONS Explorer site (<u>microns-explorer.org</u>). From the site, users can browse through the large-scale EM imagery and segmentation results using Neuroglancer (J. Maitin-Shepard, https://github.com/google/neuroglancer); several example visualizations are provided to get started. All data is served from publicly readable cloud buckets hosted through Amazon AWS and Google Cloud Storage.

To enable systematic analysis without downloading hundreds of gigabytes of data, users can selectively access cloud-based data programmatically through a collection of open source Python clients (Supplemental Table 1). The functional data, including calcium traces, stimuli, behavioral measures, and more, is available in a DataJoint^68^ database which can be accessed using DataJoint’s Python API (<u>datajoint.org</u>), or is available as Neurodata Without Borders (NWB)^69^ files on the Distributed Archives for Neurophysiology Data Integration (DANDI) Archive (https://dandiarchive.org/dandiset/000402). EM imagery and segmentation volumes can be selectively accessed using cloud-volume (W. Silversmith, https://github.com/seung-lab/cloud-volume), a Python API that simplifies interacting with large scale image data. Mesh files describing the shape of cells can be downloaded with either cloud-volume or through MeshParty (S. Dorkenwald et al., https://github.com/sdorkenw/MeshParty), which also provides features for convenient mesh analysis, skeletonization and visualization. These meshes can be decomposed and richly annotated for automated proofreading and morphological analysis of processes and spines using NEURD^50^ (https://github.com/reimerlab/NEURD). Annotations on the structural data, such as synapses and cell body locations, can be queried via CAVEclient, a Python interface to the Connectome Annotation Versioning Engine (CAVE) APIs (Fig 6a,b). CAVE encompasses a set of microservices for collaborative proofreading and analysis of large scale volumetric data.

**Figure 6.**
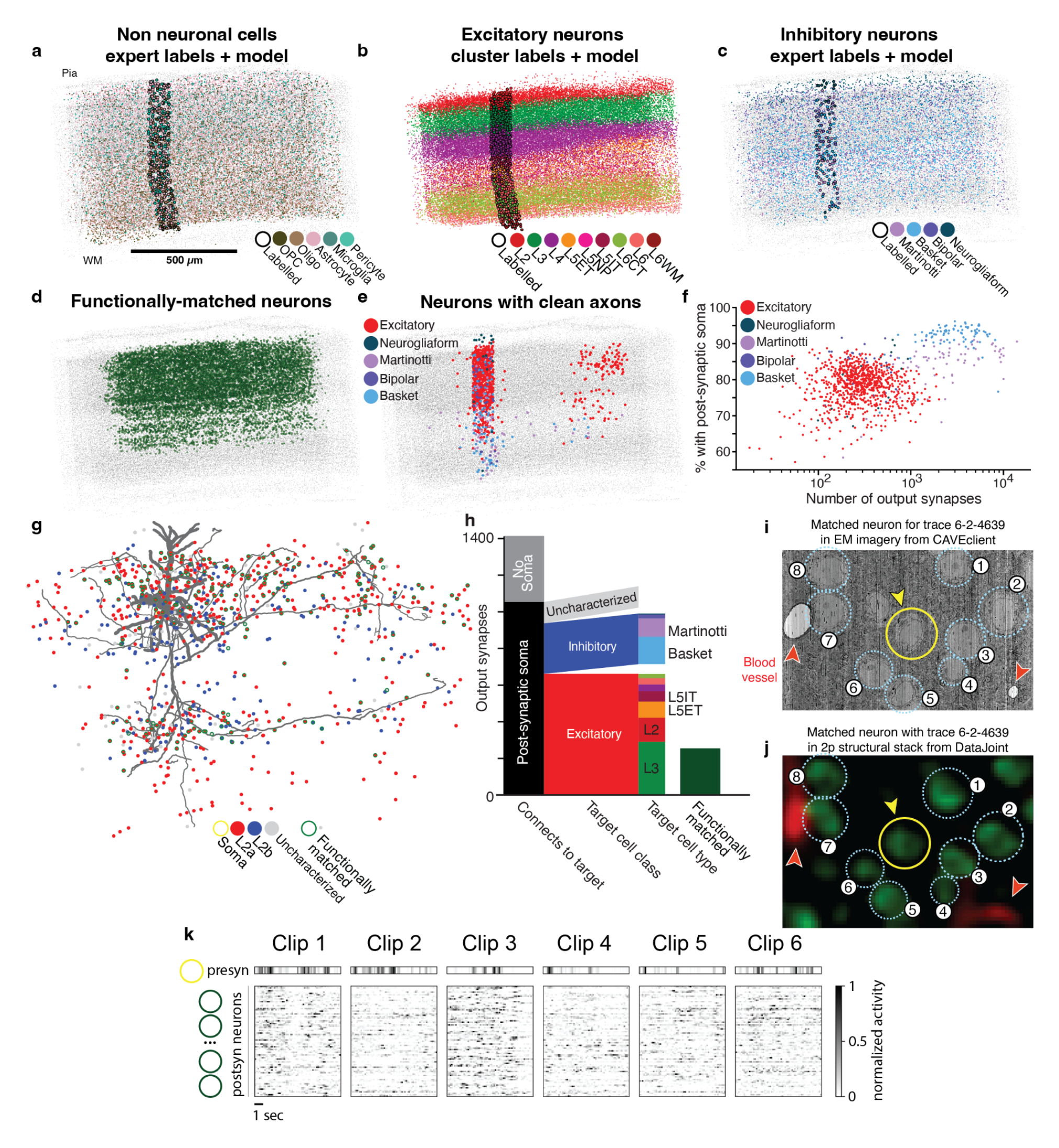
Integrated analysis resources and examples. **a-e)** An overview of the cell body locations and cell type classifications available in the dataset. All nucleus detection locations shown in light grey for spatial context. **a)** Locations of non-neuronal cells colored with respect to type, both manually typed by experts (dark outlines) and the output of a classifier^1^ (no outline). **b)** Locations of excitatory cell classes, including a labeling based on a simplified representation of an unsupervised clustering of morphological features^2^ (dark outline), and a model based on those labels^1^. **c)** Locations of inhibitory cells, including broad cell classifications by human experts^2^ and models trained on those labels^1^. **d)** The locations of neurons which were manually functional registered to functional traces in the in-vivo data. **e)** A lateral projection showing the location and proofreading status of neurons in subvolume 65 based on data in the table “proofreading_status_public_release”. Black dots are fully proofread cells, red dots indicate cells that have been cleaned of false merges but are potentially incomplete, while blue dots have had only their dendrites cleaned and extended. **f)** A quantification of reconstructions of neurons shown in e. X axis shows the number of output synapses in each reconstruction, the Y axis shows the fraction of those synapses which can be mapped to a target with a single identified post-synaptic soma. Each dot is a reconstruction colored by its broad cell class, grouping all excitatory cells together. **g)** A completely proofread pyramidal cell (nucleus id: 294657, segment id: 864691135926952148), with all of the post-synaptic soma locations of cells it connects to shown as colored dots, colored by cell type. Those cells which have a functionally co-registered region of interest are outlined in dark green. **h)** A quantification of how many synapses are associated with post-synaptic objects of different categories. First, what fraction map to a single post-synaptic soma. Second, of those that map, what fraction are excitatory or inhibitory neurons. For each inhibitory and excitatory class, what fraction of cells were classified into sub-classes based on the model outputs shown in b and c. Finally, how many synapses map to functionally co-registered cells. The cell and its synapses are viewable online (https://neuroglancer-demo.appspot.com/#!gs://microns-static-links/mm3/data_fig/6f.json). **i)** EM imagery (**j**) and corresponding imagery from the 2p structural stack centered on the cell shown in g (yellow circle). **k)** A representation of the functional response of the cell shown in g, and all its functionally coregistered post-synpatic targets. Each sub-panel represents the response to one of the ‘oracle’ clips shown in all functional scans. The heatmap reflects the average df/F trace of the presynaptic neuron (yellow) on top, and below, each row of the heatmap corresponds to a different post-synaptic target, sorted vertically based on the total synaptic strength from the pre-synaptic neuron.

The first collection of annotation tables available through CAVEclient focus on the larger subvolume of the dataset, which we refer to within the infrastructure by the nickname “Minnie65”, and which has been the current focus of proofreading and ongoing analysis (Supplemental Table 2). The largest table describes connectivity, containing all 337.3 million synapses and searchable by presynaptic id, postsynaptic id, and spatial location. In addition, there are several tables describing the soma location of key cells, predictions for which cells are different non-neuronal (Fig 6a), excitatory (Fig 6b), and inhibitory types (Fig 6c). There are also annotations which denote which cells have been functional coregistered (Fig 6d) and which cells have been proofread to different degrees of completion (Fig 6e.). In this release, the only table available for the smaller subvolume (“Minnie35”) contains synapses, as its segmentation and alignment occurred later and little proofreading, annotation or analysis has been conducted within it. We expect that continued proofreading and analysis of the data will lead to updated and additional tables for both portions of the data in future data releases.

This collection of tools and public data permits analyses that integrate questions of connectivity, morphology and functional properties of neurons. Here, we provide an example to suggest how the data might be used together. The power of the dataset lies in the fact that when an axon is proofread, it contains hundreds to over 10 thousand output synapses (Fig 6f). Furthermore, between 60-95% of those outputs can be accurately mapped onto their post-synaptic targets with a known soma location, depending on the cell-type and its spatial location in the volume (Fig 6f). This is because the segmentation is highly accurate for dendritic inputs, with a 99% input precision based on comparing proofread to non-proofread dendrites. In order to seed an analysis with an as-complete-as-possible cell, you might begin by using the proofreading table to identify a neuron with complete axons and dendrites and querying for all the synaptic inputs and outputs for the cell, in this case a L2/3 cell in VISp (Fig 6e). For this particular proofread neuron, 74.5% (1053/1412 synapses) are onto objects with a single nucleus (as determined from automated detection), with 275 synapses onto cells classified as inhibitory, 662 onto cells classified as excitatory, and 116 onto cells whose soma did not pass classification quality control (Fig 6h). The remainder (25.4% 359/1412 synapses), are onto orphan fragments, composed of a mix of disconnected spine heads and stretches of dendrite. By filtering the synaptic targets with functionally-matched neurons (Fig 6h), you can further identify which targets have been matched to the functional experiments (254/1412) and use DataJoint to query the functional data (Fig 6h–l). In this case, the targets include pyramidal cells in both L2/3 and L5. Subsequent investigation could look at the morphology of such cells in detail, or consider functional responses of their targets. We have provided example notebooks walking through the above examples and more to help users get started. Taken together, these data provide a platform for analysis of the relationship between the synaptic structure, neuronal morphology, and functional tuning of mouse visual circuits.

## Discussion

Electron microscopy is widely recognized as the gold standard for identifying synapses’ structural features, and most datasets, including the MICrONS project’s output, were primarily created to answer questions related to circuit-level connectivity. Regardless of the original intent, the scale and high resolution of the MICrONS dataset offers information that is far richer and of broader interest than just connectivity. For example, the imagery also reveals the intracellular machinery of cells, including the morphology of vital subcellular structures such as the nucleus, mitochondria, endoplasmic reticulum, and microtubules. Furthermore, the segmentation includes non-neuronal cells such as microglia, astrocytes, oligodendrocyte precursor cells and oligodendrocytes, as well the fine morphology of the cortical vasculature.

### Challenges of analyzing large scale data

The scale of large functional and EM datasets presents a wealth of opportunities for analysis and discovery. With advances in microscopy and computing power, it is now possible to work with datasets that are orders of magnitude larger than just a few years ago with millions of synapses and tens of thousands of recorded neurons. Among the key opportunities presented by this data is the ability to identify patterns and trends that may be hidden in smaller datasets, the ability to identify and validate general principles at a larger scale, and the ability to perform more sophisticated analyses – since with more data, it is possible to use more complex algorithms and models including machine learning techniques. These approaches can help to identify patterns and trends that would be difficult to observe using smaller datasets. When analyzing the connectivity graph, it’s essential to keep in mind that the automatic segmentation of dendritic inputs is highly accurate (see results). However, it’s equally important to note that the automatic segmentation of axons is not as accurate. Therefore, it’s essential to be mindful of which processes have been proofread and which have not. Additionally, it’s worth considering that although each neuron in the dataset receives thousands of inputs, a percentage of synapses in the dataset are on detached spines. Depending on the scientific question being asked, it’s worth considering if these detached spines may create bias in the conclusions drawn, such as distinguishing between excitatory and inhibitory inputs.

In the functional data, it is important to recognize that photon scattering and out-of-plane fluorescence may cause signal degradation and contamination with increasing depth from the pia surface, especially given the dense GCaMP6s expression in excitatory somas and neurites ^70, 71^. Caution should be taken to disentangle true biological variation in neuronal tuning across layers from these optical artifacts, by either matching controls at the same depth, or validating the finding with a method less prone to these artifacts (e.g. electrophysiology, or two-photon with more sparse or targeted labeling). Further, although all functional imaging was done in the same volume, it was done across several distinct imaging sessions. Technical factors as well as changes in the physiological state of the mouse should be taken into account when analyzing functional recordings taken at different times.

### Comparison with other EM connectomics studies

The importance of high resolution structural data was recognized early in invertebrate systems, particularly in the worm^72, 73^. However, it is in the fly that connectomics as the pursuit of complete connectivity diagrams has had the strongest renaissance. EM volumes now describe the *Drosophila* nervous system at both larval^74^ and adult ^42, 75^ life stages and in both central brain ^42, 75^ and nerve cord^62^. The size of volume required to capture most central neurons and their synaptic connections is well-suited to EM. The whole fly brain fills about a third of a 750 x 350 x 250 µm^3^ bounding box while the nerve cord fills about a third of a 950 x 320 x 200 µm^3^ bounding box^62^, well within the bounds of contemporary EM methods. The creation of these datasets has spurred investment in both manual skeletonized reconstruction and automated dense reconstructions ^43, 76–78^ with both centralized and community minded efforts to proofread and mine them for biological insight ^31, 79–83^. In addition to numerous targeted reconstructions in these datasets, large-scale proofread reconstructions from these datasets now include a manually traced full larval brain, a densely segmented and extensively proofread partial central brain and a publicly announced densely segmented and proofread complete adult brain. These datasets collectively span nearly the entire fly nervous system and are driving a revolution in how fly systems neuroscience is being studied.

In the mammalian system there is currently no EM dataset that contains a complete area, let alone a complete brain. There is however, as mentioned above, an established culture of making data open and publicly available ^28, 30, 31, 37, 84–86^. In the last 10 years there have only been three other rodent EM datasets which are at least 5% the size of the MICrONS multi-area dataset presented in this manuscript, with publicly available reconstructions. One dataset is 424 × 429 × 274 µm^3^ from p26 rat entorhinal cortex^87^, with skeleton reconstructions of incomplete dendrites of 667 neurons, and skeleton reconstructions of local axons of 22 excitatory neurons averaging 550 µm in length. A dataset from mouse LGN that is 500 x 400 x 280 µm^3^ is publicly available^88^, containing ∼3000 neuronal cell bodies. This dataset is large enough that dendritic reconstructions from the center of the volume are nearly complete, and it has a sparse manual segmentation, covering ∼1% of the volume, that includes 304 thalamocortical cells and 162 axon fragments. The third is a 424 × 453 × 360 µm volume covering layer 4 of mouse primary somatosensory cortex, with manual reconstruction of 52 interneuronal dendrites and numerous axons^89^.

It is critically important to compare circuit architectures across species. The neocortex is of particular interest as it is expanded in the human compared to the mouse. There is already a large body of literature on the comparative aspects between the cortex of humans and of other species. This research includes morphological and electrical properties of neurons, density of spines, synapses and neurons, as well as biophysical properties and morphology of synaptic connections ^90–96^. Importantly, a recent EM connectomics dataset of the human medial temporal gyrus^97^ vastly expands the possibilities of this comparison. This is a mm^3^ scale volume, with a maximum extent of 3 mm x 2 mm and a thickness of 150 µm. This human dataset is publicly available, including a dense automated reconstruction of all objects, with ∼16,000 neurons, 130 million synapses and an initial release of 104 proofread cells. These human connectomics data will doubtless yield critical insights. One practical difference from the volume described here is the aspect ratio of the human data, which is matched to the greater thickness of human cortex compared to mouse. To some extent the wide and thin dimensions of the human dataset trades off completeness of local neurons and circuits in order to sample all layers, while the nearly cubic volume described here is more suitable for studying local circuits and long-range connections across areas. Apart from the study by Hua and colleagues^89^, the other studies mentioned above do not have corresponding functional characterizations of the neurons reconstructed in EM. In contrast, the functional connectomics data we have released includes both anatomy and activity of the same cells.

### An opportunity to map cell types at scale

In the mammalian nervous system, transcriptomics has been the most scalable approach for cell type taxonomies. In smaller organisms like the fly, where we have both extensive gene expression maps, whole-brain neuronal reconstructions, and nearly complete connectomes, integration across modalities has been a powerful engine of discovery. Moreover, the availability of connectomes in the fly have allowed for a much higher resolution of cell types, with novel taxonomies and new cell types being discovered^41^. Our accompanying manuscripts suggest that a similar path to cell type discovery will be enabled by large-scale EM in the mammalian system with novel cell types and novel patterns of connectivity.

While this wealth of structural data on cell types and circuits provides strong constraints on the nature of the computations the brain performs, genes provide constraints on how this structure is built and operates. Linking connectomics to transcriptomics is a Frst step for merging connectivity with molecular information and building cell-type specific tools informed by how neurons connect. In one of the accompanying manuscripts we offer a proof of concept for Martinotti cells on how to achieve this link, using morphology as a common feature to integrate PatchSeq and EM datasets, suggesting a broader pathway for multimodal integration.

### The importance of functional connectomics

Almost fifty years later after Crick described his “impossible” experiment, we have provided a first draft, but its full promise will take some time to achieve. Most importantly, complete segmentation still requires an extensive amount of proofreading for the largest data sets, such as the millimeter-scale cortical reconstruction reported here. Similarly, simultaneously recording single action potentials from tens of thousands of neurons is constrained by sensor dynamics and optical sampling constraints.

Nonetheless, there has been steady progress. The first structure-function studies which combined two photon microscopy and electron microscopy examined how the wiring of mouse retina ^32–36^ and mouse visual cortex^28^ related to functional properties. Lee and colleagues^29^ related visual tuning properties of 50 functionally-characterized neurons in primary visual cortex to their connectivity measured via EM reconstruction of a 450×450×150 µm volume. 1000 synapses were mapped by hand, yielding a graph of connectivity between 29 orientation-tuned cells (a subset of the characterized cells, as in the current data set). Subsequently, our consortium used dense segmentation plus proofreading of a 250×140×90 µm dataset ^30, 31, 37^ from mouse layer 2/3 visual cortex, yielding many more overall connections, but still only twice the number of functionally characterized cells. Perhaps most impressively, In the olfactory bulb of the zebrafish, Wanner and colleagues^98^ manually reconstructed almost all neurons (n=1,003) within a 72×108×119 µm^3^ volume, in which responses to odors were measured in vivo. Their analysis of the 18,483 measured connections revealed how this structural network mediated de-correlation and variance normalization of the functional responses and demonstrates how larger measurements of network structure and function can provide mechanistic insights.

In contrast, the data released here contains tens of thousands of neurons with functionally characterized responses to visual stimuli and, because it is *densely* segmented and contains complete dendritic and local axonal arbors of centrally located cells, the opportunities to study connected neurons are orders of magnitude greater. As an example, from just 94 proofread excitatory axons, one can query 69,962 output synapses, which map to 20,112 distinct neuron soma in the volume.

Moreover, inspired by recent advancements in artificial intelligence we also created a functional digital twin of the MICrONS mouse that can enable a more comprehensive analysis of function ^7, 8^. Specifically, in an accompanying paper we developed a "foundation model" for the mouse visual cortex using deep learning which was trained using large scale data sets from multiple visual cortical areas and mice, recorded while they viewed ecological videos. The model demonstrated its generalization abilities by accurately predicting neuronal responses not only to natural videos, but also to various new stimulus domains, such as coherent moving dots and noise patterns, as confirmed through in vivo testing ^7, 8^. By applying the foundation model to the MICrONS mouse data, we created a functional digital twin of this mouse, paving the way for a systematic exploration of the relationship between circuit structure and function for tens of thousands of neurons connected with millions of synapses. Combined with the anatomical data from this mouse, we can investigate the structure-function relationships for specific visual computations ^5, 6^ and decipher the principles that determine the synaptic network in the cortex ^7, 8^. The most important goal of connectomics is to map the connections between cells, from cell body, to axon, to synapse, and back to cell body. In a large volume with complete and segmented dendrites and local axons, this can be achieved. Currently, the dendrites are nearly completely segmented (fig. 6), but many axons require proofreading. A goal in future years will be to complete the segmentation, through a combination of additional machine learning and improved proofreading. This echoes the successful strategy in the reconstruction of the Fly adult brain, which started with the TEM volume ^42^, then added the tools developed by the MICrONS program for segmentation and proofreading ^78^ and will soon release the complete connectome. If, in addition, most cell bodies have physiology with single-spike resolution ^99^, then Crick’s experimental challenge will be met. These remaining hurdles may take some time to clear, but the next steps are becoming apparent.

## Methods

### Mouse Lines

All procedures were approved by the Institutional Animal Care and Use Committee (IACUC) of Baylor College of Medicine. All results described here are from a single male mouse, age 65 days at onset of experiments, expressing GCaMP6s in excitatory neurons via SLC17a7-Cre and Ai162 heterozygous transgenic lines (recommended and generously shared by Hongkui Zeng at Allen Institute for Brain Science; JAX stock 023527 and 031562, respectively). In order to select this animal, 31 (12 female, 19 male) GCaMP6-expressing animals underwent surgery as described below. Of these, 8 animals were chosen based on a variety of criteria including surgical success and animal recovery, the accessibility of lateral higher visual areas in the cranial window, the degree of vascular occlusion, and the success of cortical tissue block extraction and staining. Of these 8 animals, one was chosen for 40 nm slicing and EM imaging based on overall quality using these criteria.

### Timeline

DOB: 12/19/17

Surgery: 2/21/18 (P64)

Two-photon imaging start: 3/4/18 (P75)

Two-photon imaging end: 3/9/18 (P80)

Structural Stack: 3/12/18 (P83)

Perfusion: 3/16/18 (P87)

### Surgery

Anesthesia was induced with 3% isoflurane and maintained with 1.5 - 2% isoflurane during the surgical procedure. Mice were injected with 5-10 mg/kg ketoprofen subcutaneously at the start of the surgery. Anesthetized mice were placed in a stereotaxic head holder (Kopf Instruments) and their body temperature was maintained at 37 °C throughout the surgery using a homeothermic blanket system (Harvard Instruments). After shaving the scalp, bupivicane (0.05 mL, 0.5%, Marcaine) was applied subcutaneously, and after 10-20 minutes an approximately 1 cm^2^ area of skin was removed above the skull and the underlying fascia was scraped and removed. The wound margins were sealed with a thin layer of surgical glue (VetBond, 3M), and a 13 mm stainless-steel washer clamped in the headbar was attached with dental cement (Dentsply Grip Cement). At this point, the mouse was removed from the stereotax and the skull was held stationary on a small platform by means of the newly attached headbar. Using a surgical drill and HP 1/2 burr, a 4 mm diameter circular craniotomy was made centered on the border between primary visual cortex and lateromedial visual cortex (V1,LM; 3.5 mm lateral of the midline, ∼1mm anterior to the lambda suture), followed by a durotomy. The exposed cortex was washed with ACSF (25 mM NaCl, 5 mM KCl, 10 mM Glucose, 10 mM HEPES, 2 mM CaCl2, 2 mM MgSO4) with 0.3 mg/mL gentamicin sulfate (Aspen Veterinary Resources). The cortical window was then sealed with a 4 mm coverslip (Warner Instruments), using cyanoacrylate glue (VetBond). The mouse was allowed to recover for 1 day prior to imaging. After imaging, the washer was released from the headbar and the mouse was returned to the home cage. Prior to surgery and throughout the imaging period, mice were singly-housed and maintained on a reverse 12-hour light cycle (off at 11 am, on at 11 pm).

### Two Photon Imaging

Mice were head-mounted above a cylindrical treadmill and calcium imaging was performed using Chameleon Ti-Sapphire laser (Coherent) tuned to 920 nm and a large field of view mesoscope (Sofroniew et al. 2016) equipped with a custom objective (excitation NA 0.6, collection NA 1.0, 21 mm focal length). Laser power after the objective was increased exponentially as a function of depth from the surface according to:

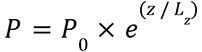

Here P is the laser power used at target depth z, P_0_ is the power used at the surface (not exceeding 10 mW), and L_z_ is the depth constant (not less than 150 μm.) Maximum laser output of 115 mW was used for scans approximately 450-500 μm from the surface and below.

### Monitor Positioning

Visual stimuli were presented to the left eye with a 31.8 x 56.5 cm (height x width) monitor (ASUS PB258Q) with a resolution of 1080 x 1920 pixels positioned 15 cm away from the eye. When the monitor is centered on and perpendicular to the surface of the eye at the closest point, this corresponds to a visual angle of ∼3.8 °/cm at the nearest point and 0.7 °/cm at the most remote corner of the monitor. As the craniotomy coverslip placement during surgery and the resulting mouse positioning relative to the objective is optimized for imaging quality and stability, uncontrolled variance in animal skull position relative to the washer used for head-mounting was compensated with tailored monitor positioning on a six dimensional monitor arm. The pitch of the monitor was kept in the vertical position for all animals, while the roll was visually matched to the roll of the animal’s head beneath the headbar by the experimenter. In order to optimize the translational monitor position for centered visual cortex stimulation with respect to the imaging field of view, we used a dot stimulus with a bright background (maximum pixel intensity) and a single dark square dot (minimum pixel intensity). Dot locations were randomly ordered from a 5 x 8 grid to tile the screen, with 15 repetitions of 200 ms presentation at each location. The final monitor position for each animal was chosen in order to center the population receptive field of the scan field ROI on the monitor, with the yaw of the monitor visually matched to be perpendicular to and 15 cm from the nearest surface of the eye at that position. A L-bracket on a six dimensional arm was fitted to the corner of the monitor at this location and locked in position, so that the monitor could be returned to the chosen position between scans and across days.

### Imaging Site Selection

The craniotomy window was leveled with regards to the objective with six degrees of freedom, five of which were locked between days to allow us to return to the same imaging site using the z-axis. Pixel-wise responses from a 3000 x 3000 μm ROI spanning the cortical window (150 μm from surface, five 600 x 3000 μm fields, 0.2 px/μm) to drifting bar stimuli were used to generate a sign map for delineating visual areas^100^. Our target imaging site was a 1200 x 1100 x 500 μm volume (anteroposterior x mediolateral x radial depth) spanning L2--L6 at the conjunction of lateral primary visual cortex (VISp) and three higher visual areas: lateromedial area (VISlm), rostrolateral area (VISrl), and anterolateral area (VISal). This resulted in an imaging volume that was roughly 50% VISp and 50% higher visual area (HVA). This target was chosen to maximize the number of visual areas within the reconstructed cortical volume, as well as maximizing the overlap in represented visual space. The imaging site was further optimized to minimize vascular occlusion and to minimize motion artifact, especially where the brain curves away from the skull/coverslip towards the lateral aspect.

Once the imaging volume was chosen, a second retinotopic mapping scan with the same stimulus was collected at 12.6 Hz and matching the imaging volume FOV with four 600 x 1100 μm fields per frame at 0.4 px/μm xy resolution to tile a 1200 x 1100 μm FOV at two depths (two planes per depth, with no overlap between coplanar fields). Area boundaries on the sign map were manually annotated.

### Two Photon Functional Imaging

Of nineteen completed scans over 6 days of imaging, fourteen are described here. Scan placement targeted 10-15 μm increments in depth to maximize coverage of the volume in depth.

- For 11 scans, imaging was performed at 6.3 Hz, collecting eight 620 x 1100 μm fields per frame at 0.4 px/μm xy resolution to tile a 1200 x 1100 μm FOV at four depths (two planes per depth, 40 μm overlap between coplanar fields).
- For 2 scans, imaging was performed at 8.6 Hz, collecting six 620 x 1100 μm fields per frame at 0.4 px/μm xy resolution to tile a 1200 x 1100 μm FOV at three depths (two planes per depth, 40 μm overlap between coplanar fields).
- For 1 scan, imaging was performed at 9.6 Hz, collecting four 620 x 1000 μm fields per frame at 0.6 px/μm xy resolution to tile a 1200 x 1000 μm FOV at two depths (two planes per depth, 40 μm overlap between coplanar fields).

In addition to locking the craniotomy window mount between days, the target imaging site was manually matched each day to preceding scans within several microns using structural features including horizontal blood vessels (which have a distinctive z-profile) and patterns of somata (identifiable by GCaMP6s exclusion as dark spots).

The full two photon imaging processing pipeline is available at (https://github.com/cajal/pipeline). Raster correction for bidirectional scanning phase row misalignment was performed by iterative greedy search at increasing resolution for the raster phase resulting in the maximum cross-correlation between odd and even rows. Motion correction for global tissue movement was performed by shifting each frame in X and Y to maximize the correlation between the cross-power spectra of a single scan frame and a template image, generated from the Gaussian-smoothed average of the Anscombe transform from the middle 2000 frames of the scan. Neurons were automatically segmented using constrained non-negative matrix factorization, then deconvolved to extract estimates of spiking activity, within the CAIMAN pipeline^57^. Cells were further selected by a classifier trained to separate somata versus artifacts based on segmented cell masks, resulting in exclusion of 8.1% of masks.

### Two Photon Structural Stack

Approximately 55 minutes prior to collecting the stack, the animal was injected subcutaneously with 60 μL of 8.3 mM Dextran Texas Red fluorescent dye (Invitrogen, D3329). The stack was composed of 30 repeats of three 1300 x 620 μm fields per depth in two channels (green and red, respectively), tiling a 1300 x 1400 μm field of view (460 μm total overlap) at 335 depths from 21 μm above the surface to 649 μm below the surface. The green channel average image across repetitions for each field was enhanced with local contrast normalization using a gaussian filter to calculate the local pixel means and standard deviations. The resulting image was then gaussian smoothed and sharpened using a Laplacian filter. Enhanced and sharpened fields were independently stitched at each depth. The resulting stitched planes were independently horizontally and vertically aligned by maximizing the correlation of the cross-power spectrum of their Fourier transformations. Finally, the resulting alignment was detrended in Z using a Hann filter with a size of 60 μm to remove the influence of vessels passing through the fields. The resulting transform was applied to the original average images resulting in a structural two photon 1322 x 1412 x 670 μm volume at 0.5 x 0.5 x 0.5 px/μm resolution in both red and green channels.

Due to tissue deformation from day to day across such a wide field of view, some cells are recorded in more than one scan. To assure we count cells only once, we subsample our recorded cells based on proximity in 3-d space. Functional scan fields were independently registered using an affine transformation matrix with 9 parameters estimated via gradient ascent on the correlation between the sharpened average scanning plane and the extracted plane from the sharpened stack. Using the 3-d centroids of all segmented cells, we iteratively group the closest two cells from different scans until all pairs of cells are at least 10 μm apart or a further join produces an unrealistically tall mask (20 μm in z). Sequential registration of sections of each functional scan into the structural stack was performed to assess the level of drift in the z dimension. All scans had less than 10 μm drift over the 1.5 hour recording, and for most of them drift was limited to <5 μm.

Fields from the FOV matched retinotopy scan described above were registered into the stack using the same approach, and the manually annotated area masks were transformed into the stack. These area masks were extended vertically across all depths, and functional units inherit their area membership from their stack xy coordinates.

### Eye and Face Camera

Movie of the animal’s eye and face was captured throughout the experiment. A hot mirror (Thorlabs FM02) positioned between the animal’s left eye and the stimulus monitor was used to reflect an IR image onto a camera (Genie Nano C1920M, Teledyne Dalsa) without obscuring the visual stimulus. An infrared 940 nm LED (Thorlabs M940L2) illuminated the right side of the animal, backlighting the silhouette of the face. The position of the mirror and camera were manually calibrated per session and focused on the pupil. Field of view was manually cropped for each session (ranging from 828 x 1217 pixels to 1080 x 1920 pixels at ∼20 Hz), such that the field of view contained the superior, frontal, and inferior portions of the facial silhouette as well as the left eye in its entirety. Frame times were time stamped in the behavioral clock for alignment to the stimulus and scan frame times. Video was compressed using Labview’s MJPEG codec with quality constant of 600 and stored the frames in AVI file.

Light diffusing from the laser during scanning through the pupil was used to capture pupil diameter and eye movements. Notably, scans using wide ranges in laser power to scan both superficial and deep planes resulted in a variable pupil intensity between frames. A custom semi-automated user interface in Python was built for dynamic adaptation of fitting parameters throughout the scan to maximize pupil tracking accuracy and coverage. The video is manually cropped to a rectangular region that includes the entirety of the eye at all time points. The video is further manually masked to exclude high intensity regions in the surrounding eyelids and fur. In cases where a whisker is present and occluding the pupil at some time points, a merge mask is drawn to bridge ROIs drawn on both sides of the whisker into a single ROI. For each frame, the original and filtered image is visible to the user. The filtered image is an exponentially-weighted temporal running average, which undergoes exponentiation, gaussian blur, automatic Otsu thresholding into a binary image, and finally pixel-wise erosion / dilation. In cases where only one ROI is present, the contour of the binary ROI is fit with an ellipse by minimizing least squares error, and for ellipses greater than the minimum contour length the xy center and major and minor radii are stored. In cases where more than one ROI is present, the tracking is automatically halted until the user either resolves the ambiguity, or the frame is not tracked (a NaN is stored). Processing parameters were under dynamic control of the user, with instructions to use the minimally sufficient parameters that result in reliably and continuous tracing of the pupil, as evidenced by plotting of the fitted ROI over the original image. Users could also return to previous points in the trace for re-tracking with modified processing parameters, as well as manually exclude periods of the trace in which insufficient reliable pupil boundary was visible for tracking.

### Treadmill

The mouse was head-restrained during imaging but could walk on a treadmill. Rostro-caudal treadmill movement was measured using a rotary optical encoder (Accu-Coder 15T-01SF-2000NV1ROC-F03-S1) with a resolution of 8000 pulses per revolution, and was recorded at ∼57-100 Hz in order to extract locomotion velocity.

### Stimulus Composition

Each scan stimulus is approximately 84 minutes and comprised:

- **Oracle Natural Movies:** 6 natural movie clips, 2 from each category. 10 seconds each, 10 repeats per scan, 10 minutes total. Conserved across all scans.
- **Unique Natural Movies:** 144 natural movies, 48 from each category. 10 seconds each, 1 repeat per scan, 24 minutes total. Unique to each scan.
- **2x Repeat Natural Movies:** 90 natural movies, 30 from each category. 10 seconds each, 2 repeats (one in each half of the scan), 30 minutes total. Conserved across all scans.
- **Local Directional Parametric Stimulus ("Trippy"):** 20 seeds, 15 seconds each, 2 repeats (one in each half of the scan), 10 minutes total. 10 seeds conserved across all scans, 10 unique to each scan.
- **Global Directional Parametric Stimulus ("Monet"):** 20 seeds, 15 seconds each, 2 repeats (one in each half of the scan), 10 minutes total. 10 seeds conserved across all scans, 10 unique to each scan.

Each scan is also preceded by 0.15 - 5.5 minutes with the monitor on, and followed by 8.3 - 21.2 minutes with the monitor off, in order to collect spontaneous neural activity.

### Natural Visual Stimulus

The visual stimulus was composed of dynamic stimuli, primarily including natural video but also including generated parametric stimuli with strong local or global directional component. Natural video clips were 10 second clips from one of three categories:

- **Cinematic** from the following sources -- Mad Max: Fury Road (2015), Star Wars: Episode VII - The Force Awakens (2015), The Matrix (1999), The Matrix Reloaded (2003), The Matrix Revolutions (2003), Koyaanisqatsi: Life Out of Balance (1982), Powaqqatsi: Life in Transformation (1988), Naqoyqatsi: Life as War (2002).
- **Sports-1M** collection (Karpathy et al. 2014) with the following keywords: cycling, mountain unicycling, bicycle, bmx, cyclo-cross, cross-country cycling, road bicycle racing, downhill mountain biking, freeride, dirt jumping, slopestyle, skiing, skijoring, alpine skiing, freestyle skiing, greco-roman wrestling, luge, canyoning, adventure racing, streetluge, riverboarding, snowboarding, mountainboarding, aggressive inline skating, carting, freestyle motocross, f1 powerboat racing, basketball, base jumping.
- **Rendered 3D video** of first person POV random exploration of a virtual environment with moving objects, produced in a customized version of Unreal Engine 4 with modifications that enable precise control and logging of frame timing and camera positions to ensure repeatability across multiple rendering runs. Environments and assets were purchased from Unreal Engine Marketplace. Assets chosen for diversity of appearance were translated along a piecewise linear trajectory, and rotated with a piece-wise constant angular velocity. Intervals between change points were drawn from a uniform distribution from 1 to 5 seconds. If a moving object encountered an environmental object, it bounced off and continued along a linear trajectory reflected across the surface normal. The first person POV camera followed the same trajectory process as the moving objects. Light sources were the default for the environment. Latent variable images were generated by re-generating the scenes and trajectories, rendering different properties, including: absolute depth, object identification number, and surface normals.

All natural movies were temporally resampled to 30 fps, and were converted to grayscale with 256 x 144 pixel resolution with FFmpeg (ibx264 at YUV4:2:0 8bit). Stimuli were automatically filtered for upper 50th percentile Lucas-Kanade optical flow and temporal contrast of the central region of each clip. All natural movies included in these experiments were further manually screened for unsuitable characteristics (ex. fragments of rendered videos in which the first person perspective would enter a corner and become "trapped" or follow an unnatural camera trajectory, or fragments of cinematic or Sports-1M containing screen text or other post-processing editing).

### Global Directional Parametric Stimulus ("Monet")

To probe neuronal tuning to orientation and direction of motion, a visual stimulus was designed in the form of smoothened Gaussian noise with coherent orientation and motion. Briefly, an independently identically distributed (i.i.d.) Gaussian noise movie was passed through a temporal low-pass Hamming filter (4 Hz) and a 2-d Gaussian spatial filter (σ = 3.0° at the nearest point on the monitor to the mouse). Each 15-second block consisted of 16 equal periods of motion along one of 16 unique directions of motion between 0-360 degrees with a velocity of 42.8 degrees/s at the nearest point on the monitor. The movie was spatial filtered to introduce a spatial orientation bias perpendicular to the direction of movement by applying a bandpass Hanning filter G(ω; c) in the polar coordinates in the frequency domain for ω = ϕ − θ where ϕ is the polar angle coordinate and θ is the movement direction θ. Then:

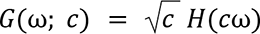

and

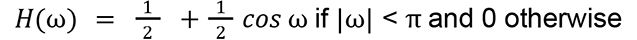

Here, c=2.5 is an orientation selectivity coefficient. At this value, the resulting orientation kernel’s size is 72° full width at half maximum in spatial coordinates.

### Local Directional Parametric Stimulus ("Trippy")

To probe the tuning of neurons to local spatial features including orientation, direction, spatial and temporal frequency, the "Trippy" stimulus was synthesized by applying the cosine function to a smoothened noise movie. Briefly, a *phase movie* was generated as an independently identically distributed (i.i.d.) uniform noise movie with 4 Hz temporal bandwidth. The movie was upsampled to 60 Hz with the Hanning temporal kernel. An increasing trend of 8π/*s* was added to the movie to produce drifting grating movements whereas the noise component added local variations of the spatial features. The movie was spatially upsampled to the full screen with a 2D Gaussian kernel with a sigma of 5.97 cm or 22.5° at the nearest point. The resulting stimulus yielded the local phase movie of the gratings, from which all visual features are derived analytically.

### Stimulus Alignment

A photodiode (TAOS TSL253) was sealed to the top left corner of the monitor, where stimulus sequence information was encoded in a 3 level signal according to the binary encoding of the flip number assigned in-order. This signal was recorded at 10 MHz on the behavior clock (MasterClock PCIe-OSC-HSO-2 card). The signal underwent a sine convolution, allowing for local peak detection to recover the binary signal. The encoded binary signal was reconstructed for 89% of trials. A linear fit was applied to the trial timestamps in the behavioral and stimulus clocks, and the offset of that fit was applied to the data to align the two clocks, allowing linear interpolation between them.

### Oracle Score

We used six natural movie conditions that were present in all scans and repeated 10 times per scan to calculate an "Oracle score" representing the reliability of the trace response to repeated visual stimuli. This score was computed as the jackknife mean of correlations between the leave-one-out mean across repeated stimuli with the remaining trial.

### Tissue Preparation

After optical imaging at Baylor College of Medicine, candidate mice were shipped via overnight air freight to the Allen Institute. All procedures were carried out in accordance with the Institutional Animal Care and Use Committee at the Allen Institute for Brain Science. All mice were housed in individually ventilated cages, 20-26 C, 30-70% Relative Humidity, with a 12-hour light/dark cycle. Mice were transcardially perfused with a fixative mixture of 2.5% paraformaldehyde, 1.25% glutaraldehyde, and 2 mM calcium chloride, in 0.08 M sodium cacodylate buffer, pH 7.4. After dissection, the neurophysiological recording site was identified by mapping the brain surface vasculature. A thick (1200 um) slice was cut with a vibratome and post-fixed in perfusate solution for 12 – 48 h. Slices were extensively washed and prepared for reduced osmium treatment (rOTO) based on the protocol of Hua and colleagues ^101^. All steps were performed at room temperature, unless indicated otherwise. 2% osmium tetroxide (78 mM) with 8% v/v formamide (1.77 M) in 0.1 M sodium cacodylate buffer, pH 7.4, for 180 minutes, was the first osmication step. Potassium ferricyanide 2.5% (76 mM) in 0.1 M sodium cacodylate, 90 minutes, was then used to reduce the osmium. The second osmium step was at a concentration of 2% in 0.1 M sodium cacodylate, for 150 minutes. Samples were washed with water, then immersed in thiocarbohydrazide (TCH) for further intensification of the staining (1% TCH (94 mM) in water, 40 °C, for 50 minutes). After washing with water, samples were immersed in a third osmium immersion of 2% in water for 90 minutes. After extensive washing in water, lead aspartate (Walton’s (20 mM lead nitrate in 30 mM aspartate buffer, pH 5.5), 50 °C, 120 minutes) was used to enhance contrast. After two rounds of water wash steps, samples proceeded through a graded ethanol dehydration series (50%, 70%, 90% w/v in water, 30 minutes each at 4 °C, then 3 x 100%, 30 minutes each at room temperature). Two rounds of 100% acetonitrile (30 minutes each) served as a transitional solvent step before proceeding to epoxy resin (EMS Hard Plus). A progressive resin infiltration series (1:2 resin:acetonitrile (e.g. 33% v/v), 1:1 resin:acetonitrile (50% v/v), 2:1 resin acetonitrile (66% v/v), then 2 x 100% resin, each step for 24 hours or more, on a gyrotary shaker) was done before final embedding in 100% resin in small coffin molds. Epoxy was cured at 60 °C for 96 hours before unmolding and mounting on microtome sample stubs for trimming.

The surface of the brain in the neurophysiology ROI was highly irregular, with depressions and elevations that made it impossible to trim all the resin from the surface of the cortex without removing layer 1 (L1) and some portions of layer 2 (L2). Though empty resin increases the number of folds in resulting sections, we left some resin so as to keep the upper layers (L1 and L2) intact to preserve inter-areal connectivity and the apical tufts of pyramidal neurons. Similarly, white matter was also maintained in the block to preserve inter-areal connections despite the risk of increased sectioning artifacts that then have to be corrected through proofreading.

### Ultrathin Sectioning

The sections were then collected at a nominal thickness of 40 nm using a modified ATUMtome^61^ (RMC/Boeckeler) onto 6 reels of grid tape. The knife was cleaned every 100–500 sections, occasionally leading to the loss of a very thin partial section (≪ 40 nm). Thermal expansion of the block as sectioning resumed post-cleaning resulted in a short series of sections substantially thicker than the nominal cutting thickness. The sectioning took place in two sessions, the first session took 8 consecutive days on a 24/7 schedule and contained sections 1 to 14,773. The loss rate on this initial session was low, but before section 7931 there were two events that led to consecutive section loss. At the end of this session we started seeing differential compression between the resin and the surface of the cortex. Because this could lead to severe section artifacts, we paused to trim additional empty resin from the block and also replaced the knife. The second session lasted five consecutive days and an additional 13,199 sections were cut. Due to the interruption, block shape changes and knife replacement, there are approximately 45 partial sections at the start of this session; importantly, these do not represent tissue loss (see stitching and alignment section). As will be described later, the EM dataset is subdivided into two subvolumes due to sectioning and imaging events that resulted in loss of a series of sections.

### Transmission Electron Microscopy Imaging

The parallel imaging pipeline described here^61^ converts a fleet of transmission electron microscopes into high-throughput automated image systems capable of 24/7 continuous operation. It is built upon a standard JEOL 1200EXII 120kV TEM that has been modified with customized hardware and software. The key hardware modifications include an extended column and a custom electron-sensitive scintillator. A single large-format CMOS camera outfitted with a low distortion lens is used to grab image frames at an average speed of 100 ms. The autoTEM is also equipped with a nano-positioning sample stage that offers fast, high-fidelity montaging of large tissue sections and an advanced reel-to-reel tape translation system that accurately locates each section using index barcodes for random access on the GridTape. In order for the autoTEM system to control the state of the microscope without human intervention and ensure consistent data quality, we also developed customized software infrastructure piTEAM that provides a convenient GUI-based operating system for image acquisition, TEM image database, real-time image processing and quality control, and closed-loop feedback for error detection and system protection etc. During imaging, the reel-to-reel GridStage moves the tape and locates targeting aperture through its barcode. The 2D montage is then acquired through raster scanning the ROI area of tissue. Images along with metadata files are transferred to the data storage server. We perform image QC on all the data and reimage sections that fail the screening. Pixel sizes for all systems were calibrated within the range between 3.95∼4.05 nm/pix and the montages had a typical size of 1.2 mm × 0.82 mm. The EM dataset contains raw tile images with two different sizes because two cameras with two different resolutions were used during acquisition. The most commonly used was a 20MP camera that required 5,000 individual tiles to capture the 1 mm^2^ montage of each section. During the dataset acquisition, three autoTEMs were upgraded with 50MP camera sensors, which increased the frame size and reduced the total number of tiles required per montage to ∼2,600

### Volume Assembly

The images in the serial section are first corrected for lens distortion effects. A non-linear transformation of higher order is computed for each section using a set of 10 x 10 highly overlapping images collected at regular intervals during imaging^63^. The lens distortion correction transformations should represent the dynamic distortion effects from the TEM lens system and hence require an acquisition of highly overlapping calibration montages at regular intervals. Overlapping image pairs are identified within each section and point correspondences are extracted for every pair using a feature based approach. In our stitching and alignment pipeline, we use SIFT feature descriptors to identify and extract these point correspondences. Per image transformation parameters are estimated by a regularized solver algorithm. The algorithm minimizes the sum of squared distances between the point correspondences between these tile images. Deforming the tiles within a section based on these transformations results in a seamless registration of the section. A downsampled version of these stitched sections are produced for estimating a per-section transformation that roughly aligns these sections in 3D. A process similar to 2D stitching is followed here, where the point correspondences are computed between pairs of sections that are within a desired distance in z direction. The per-section transformation is then applied to all the tile images within the section to obtain a rough aligned volume. MIPmaps are utilized throughout the stitching process for faster processing without compromise in stitching quality.

The rough aligned volume is rendered to disk for further fine alignment. The software tools used to stitch and align the dataset is available in our github repository https://github.com/AllenInstitute/asap-modules. The volume assembly process is entirely based on image meta-data and transformations manipulations and is supported by the Render service (https://github.com/saalfeldlab/render).

Cracks larger than 30 um in 34 sections were corrected by manually defining transforms. The smaller and more numerous cracks and folds in the dataset were automatically identified using convolutional networks trained on manually labeled samples using 64 x 64 x 40 nm^3^ resolution image. The same was done to identify voxels which were considered tissue. The rough alignment was iteratively refined in a coarse-to-fine hierarchy^102^, using an approach based on a convolutional network to estimate displacements between a pair of images ^103^. Displacement fields were estimated between pairs of neighboring sections, then combined to produce a final displacement field for each image to further transform the image stack. Alignment was first refined using 1024 x 1024 x 40 nm^3^ images, then 64 x 64 x 40 nm^3^ images.

The composite image of the partial sections was created using the tissue mask previously computed. Pixels in a partial section which were not included in the tissue mask were set to the value of the nearest pixel in a higher-indexed section that was considered tissue. This composite image was used for downstream processing, but not included with the released images.

### Segmentation

Remaining misalignments were detected by cross-correlating patches of image in the same location between two sections, after transforming into the frequency domain and applying a high-pass filter. Combining with the tissue map previously computed, a mask was generated that sets the output of later processing steps to zero in locations with poor alignment. This is called the segmentation output mask.

Using the method outlined in^64^, a convolutional network was trained to estimate inter-voxel affinities that represent the potential for neuronal boundaries between adjacent image voxels. A convolutional network was also trained to perform a semantic segmentation of the image for neurite classifications, including (1) soma+nucleus, (2) axon, (3) dendrite, (4) glia, and (5) blood vessel. Following the methods described in (Wu et al. 2021), both networks were applied to the entire dataset at 8 x 8 x 40 nm^3^ in overlapping chunks to produce a consistent prediction of the affinity and neurite classification maps. The segmentation output mask was applied to the predictions.

The affinity map was processed with a distributed watershed and clustering algorithm to produce an over-segmented image, where the watershed domains are agglomerated using single-linkage clustering with size thresholds^65, 104^. The over-segmentation was then processed by a distributed mean affinity clustering algorithm^65, 104^ to create the final segmentation. We augmented the standard mean affinity criterion with constraints based on segment sizes and neurite classification maps during the agglomeration process to prevent neuron-glia mergers as well as axon-dendrite and axon-soma mergers.

### Synapse detection & assignment

A convolutional network was trained to predict whether a given voxel participated in a synaptic cleft. Inference on the entire dataset was processed using the methods described in^66^ (using 8 x 8 x 40 nm^3^ images). These synaptic cleft predictions were segmented using connected components, and components smaller than 40 voxels were removed.

A separate network was trained to perform synaptic partner assignment by predicting the voxels of the synaptic partners given the synaptic cleft as an attentional signal ^105^. This assignment network was run for each detected cleft, and coordinates of both the presynaptic and postsynaptic partner predictions were logged along with each cleft prediction.

### Nucleus detection

A convolutional network was trained to predict whether a voxel participated in a cell nucleus. Following the methods described in^66^, a nucleus prediction map was produced on the entire dataset at 64 x 64 x 40 nm^3^. The nucleus prediction was thresholded at 0.5, and segmented using connected components.

### Proofreading

Extensive manual, semi-automated, and fully automated proofreading of the segmentation data was performed by multiple teams to improve the accuracy of the neural circuit reconstruction. Critical to enabling these coordinated proofreading activities is the central ChunkedGraph system^31^, which maintains a dynamic segmentation dataset, and supports real-time collaborative proofreading on petascale datasets though scalable software interfaces to receive edit requests from various proofreading platforms and support querying and analysis on edit history.

Multiple proofreading platforms and interfaces were developed and leveraged to support the large-scale proofreading activities performed by various teams at Princeton University, the Allen Institute for Brain Science, Baylor College of Medicine, the Johns Hopkins University Applied Physics Laboratory, and Ariadne (see acknowledgements for complete list of individual proofreaders). Below we outline the methods for these major proofreading activities focused on improving the completeness of neurons within and proximal to the main cortical column, splitting of merged multi-soma objects distributed throughout the image volume, and distributed application of automated proofreading edits to split erroneously merged neuron segments.

#### Manual proofreading of dendritic and axonal processes

Following the methods described previously^31, 78^ proofreaders from Princeton University, the Allen Institute for Brain Science, Baylor College of Medicine, and Ariadne used a modified version of Neuroglancer with annotation capabilities as a user interface to make manual split and merge edits to neurons with somata spatially located throughout the dataset. The choice of which neurons to proofread was based on the scientific needs of different projects, which are described in accompanying manuscripts^2, 4, 7^.

Proofreading was aided by on-demand highlighting of branch points and tips on user-defined regions of a neuron based on rapid skeletonization (https://github.com/AllenInstitute/Guidebook). This approach quickly directed proofreader attention to potential false merges and locations for extension, as well as allowed a clear record of regions of an arbor that had been evaluated.

For dendrites, we checked all branch points for correctness and all tips to see if they could be extended. False merges of simple axon fragments onto dendrites were often not corrected in the raw data, since they could be computationally filtered for analysis after skeletonization (see next section). Detached spine heads were not comprehensively proofread. Dendrites that were proofread are identified in CAVE table “proofreading_status_public_release” as “clean”

For axons, we began by "cleaning" axons of false merges by looking at all branch points. We then performed an extension of axonal tips, the degree of this extension depended on the scientific goals of the different project. The different proofreading strategies were as follow:

1. Comprehensive extension: Each axon end and branch point was visited and checked to see if it was possible to extend until either their biological completion or reached an incomplete end (incomplete ends were due to either the axon reaching the borders of the volume or an artifact that curtailed its continuation).
2. Substantial extension: Each axon branch point was visited and checked, many but not all ends were visited and many but not all ends were done.
3. Inter_areal_extension: A subset of axons that projected either to an HVA from V1, or from V1 to an HVA were preferentially extended to look specifically at inter-areal connections.
4. Local L23 cylinder cutting: A subset layer 2/3 pyramidal cells were proofread in a local cylinder which had a 300 micron radius centered around the column featured in CSM et al., with a floor at the layer 4/5 border. Any axon leaving that cylinder was cut and not followed further.
5. Local L4 cylinder cutting: A subset layer 4 pyramidal cells were proofread in a local cylinder which had a 300 micron radius centered around the column featured in CSM et al., with a floor at the layer 5/6 border. Any axon leaving that cylinder was cut and not followed further.
6. At least 100 synapses: Axons were extended until at least 100 synapses were present on the axon to get a sampling of their output connectivity profile.

Axons that were proofread are identified in CAVE table “proofreading_status_public” as “clean” and the proofreading strategy associated with each axon is described in the CAVE table “proofreading_strategy”.

#### Manual Proofreading to Split Incorrectly Merged Cells

Proofreading was also performed to correctively split multi-soma objects containing more than one neuronal soma, which had been incorrectly merged from the agglomeration step in the reconstruction process. This proofreading was performed by the Johns Hopkins University Applied Physics Laboratory, Princeton University, the Allen Institute for Brain Science, and Baylor College of Medicine. These erroneously merged multi-soma objects were specifically targeted given their number, distribution throughout the volume, and subsequent impact on global neural connectivity^106^ (Supplemental Fig 3). As an example, multi-soma objects comprised up to 20% of the synaptic targets for 78 excitatory cells that with proofreading status “comprehensive extension”. While the majority of multi-soma objects contained 2 to 25 nuclei (Supplemental Fig 3a), one large multi-soma object contained 172 neuronal nuclei due to proximity to a major blood vessel present in a substantial portion of the image volume.

Different Neuroglancer web based applications ^31, 106106^ were used to perform this proofreading, but most edits were performed using NeuVue^106^. NeuVue enables scalable task management across dozens of concurrent users, as well as provide efficient queuing, review, and execution of proofreading edits by integrating with primary data management APIs such as CAVE and PCG. Multi-soma objects used to generate proofreading tasks were originally identified using the nucleus detection table available through CAVE ^1^. Additionally, algorithms were employed in a semi-automated workflow to detect the presence of incorrect merges and proposed potential corrective split locations in the segmentation for proofreaders to review and apply ^50^.

#### Proofreading through Automated Error Detection and Correction Framework

Following the methods described elsewhere ^50^ automated error detection and error correction methods were employed using the Neural De-composition (NEURD) framework to apply edits to split incorrectly merged axonal and dendritic segments distributed across the image volume. These automated methods leveraged graph filter and graph analysis algorithms to accurately identify errors in the reconstruction and generate corrective solutions. Validation and refinement of these methods was performed through manual review of proposed automated edits through the NeuVue platform ^106^.

### Co-registration

#### Transform

We initially manually matched 2934 fiducials between the EM volume and the two-photon structural dataset (1994 somata and 942 blood vessels, mostly branch points, which are available as part of the resource). Though the fiducials cover the total volume of the dataset it is worth noting that below 400 µm from the surface there is much lower signal to noise in the 2P structural dataset requiring more effort to identify somata, therefore we made use of more vascular fiducials. Using the fiducials, a transform between the EM dataset and the two-photon structural stack was calculated (see Methods). To evaluate the error of the transform we evaluated the distance in microns between the location of a fiducial after co-registration and its original location; a perfect co-registration would have residuals of zero microns). The average residual was 3.8 microns.

The fiducial annotation was done using a down-sampled EM dataset with pixel sizes 256 nm (x), 256 nm (y), 940 nm (z). Though the fiducials cover the total volume of the dataset it is worth noting that below 400 µm from the surface there is much lower signal to noise in the 2P structural dataset requiring more effort to identify somata, therefore we made use of more vascular fiducials.

For calculating the transform we introduced a staged approach to separate the gross transformation between the EM volume and the two photon space from the finer non-linear deformations needed to get good residuals (a residual is the distance in microns between the location of a fiducial after co-registration and its original location; a perfect co-registration would have residuals of zero microns). This was done by taking advantage of the infrastructure created for the alignment of the EM dataset described above.

The full 3D transform is a list of eight transforms that fall into 4 groups with different purposes:

1. The first group is a single transform that is a 2nd-order polynomial transform between the two datasets. This first group serves to scale and rotate the optical dataset into EM space, followed by a single global nonlinear term, leaving an average residual of ∼10 µm.
2. The second group of transforms addresses an issue we saw in the residuals: there were systematic trends in the residual, both positive and negative, that aligned well with the EM z-axis. These trends are spaced in a way that is indicative of changing shape of the EM data on approximately the length scale between knife-cleanings or tape changes. We addressed this with a transform that binned the data into z-ranges and applied a further 2nd-order polynomial to each bin. We did this in a 2-step hierarchical fashion, first with 5 z-bins, followed by a second with 21 z-bins. These steps removed the systematic trends in the residuals vs z and the average residuals dropped to 5.6µm and 4.6µm respectively.
3. The third group is a set of hierarchical thin plate spline transforms. We used successively finer grids of control points of even “n x n x n” spacing in the volume. We used four steps with n = [3, 5, 10, 12]. The idea here is to account for deformations on larger length scales first, so that the highest order transforms introduce smaller changes in position. The average residuals in these steps were 3.9, 3.5, 3.1, and 2.9µm accomplished with average control point motions of 12.5, 7.5, 3.8, and 1.6µm.
4. The final group is a single thin plate spline transform. The control points for this transform are no longer an evenly spaced grid. Instead, each fiducial point is assigned to be a control point. This transform minimizes the residuals almost perfectly (as it should for the control points which are identical to the fiducials; 0.003µm on average, Fig 3) and accomplishes this final step by moving each data point on average another 2.9µm. This last transform is very sensitive to error in fiducial location but provides the co-registration with minimal residuals. This last transform is also more likely to create errors in regions with strong distortions, as for example the edges of the dataset.

Since the nature of transform 4 is to effectively set the residuals to zero for the control points, we used a new measure to evaluate the error of the transform. We created 2933 3D transforms, each time leaving out one fiducial and then evaluated the residual of the left-out point. We call this measure “leave-one-out” residuals and it evaluates how well the transform does with a new point.

#### Manual Matching

A custom user interface was used to visualize images from both the functional data and EM data side-by-side to manually associate functional units to their matching EM cell counterpart and vice versa. To visualize the functional scans, summary images were generated by averaging the scan over time (average image) and correlating pixels with neighbor pixels over time (correlation image). The product of the average and correlation images were used to clearly visualize cell body locations. Using the per field affine registration into the stack, a representative image of labeled vasculature corresponding to the registered field was extracted from the stack red channel. EM imagery and EM nucleus segmentation was resized to 1 um3 resolution, and transformed into the two-photon structural stack coordinates using the coregistration, allowing an image corresponding to the registered field to be extracted. The overlay of the extracted vessel field and extracted EM image were used to confirm local alignment of the vasculature visible in both domains. Soma identity was assessed by comparing the spatial structure of the target soma and nearby somas in the functional image to soma locations from the EM cell nuclei image. Using the tool, matchers generated a hypothesis for which EM cell nucleus matched to a given functional unit or vice versa. A custom version of Neuroglancer (Seung lab, https://github.com/seung-lab/neuroglancer) was used to visualize the region of interest in the ground-truth EM data for match confirmation.

#### Automated Matching

The spline-based co-registration transform was fine-tuned with the vasculature data from the two-photon structural stack to programmatically find matches for excitatory cells. Fine tuning is performed via symmetric diffeomorphic registration on the blood vessels in a 3D registration. This is a non-rigid transformation approach that corrects for extreme nonlinearities in the transform. The input to this method is the output of the spline-based co-registration. The EM segmentation and the two-photon structural stack volumes were each subsampled to matrices of the same size with 1 μm voxels, such that the relative scaling and indexing are equivalent for both volumes.

At this point, signal pre-processing is performed to account for the inconsistent signal quality of the two-photon stack, as neurons deep to the cortical surface emit less fluorescent intensity. To address this problem, a Meijering neurite filter^107^ was utilized. This is a ridge operator designed specifically to detect neurites in fluorescence imaging through the eigenvectors of the Hessian matrix of image intensities.

An additional filtering step is taken to account for the difference in Z-resolution between the two-photon structural stack and the EM. In the two-photon structural stack, a ‘smearing’ in the Z-direction was observed due to the extreme anisotropy in the Z-direction. To account for this, the filtered two-photon structural stack and EM are binarized via threshold and skeletonized. The TEASAR-based skeletonization algorithm comes from the Kimimaro^108^ package from Seung Lab which takes a 3D segmentation and returns a one-pixel-thick spine. A Gaussian filter is convolved over this spine to create a tube of constant radius to co-register.

The local diffeomorphic transformation is computed using the DIPY symmetrical diffeomorphic registration algorithm where one volume is selected as the “static” volume and the other volume is therefore the “moving” volume^109^. The output of the symmetrical diffeomorphic registration is a flow field that dictates how each voxel in the input space maps onto the output space. The entire volume was registered at once at full resolution. An inverse matrix is calculated in parallel to determine flow from two-photon structural stack to EM.

The fine registration itself is mediated by an expectation maximization metric between the above Gaussian filtered skeletons that were thresholded to become binary segmentations. For the final matching, the flow field generated by the registration was applied to the nuclear segmentation of the EM and conversely to the centroids of the 2P units. Linear sum assignment was performed on the two lists of centroids to perform the assignment. The nuclear segmentation is filtered by only considering cells that are known to be neurons in the CAVE table nucleus_ref_neuron_svm^1^. To improve confidence, a threshold was imposed on both the residual and separation of the match. Separation is defined as the residual to the closest centroid that was not assigned via linear sum assignment minus the residual to the assigned centroid in the same coordinate space. A threshold on the residuals of 15 μm and a threshold on the separation of 5 μm was imposed to filter out low-confidence matches, yielding an agreement of 89.9% with manual matchers across 6088 nuclei (Supplemental Fig 5).

## Cell Classification

We analyzed the nucleus segmentations for features such as volume, surface area, fraction of membrane within folds, and depth in cortex. We trained an SVM machine classifier to use these features to detect which nucleus detections were likely neurons within the volume, with 96.9% precision and 99.6% recall. This model was trained based upon data from an independent dataset, and the performance numbers are based upon evaluating the concordance of the model with the manual cell type calls within the volume. This model predicted 82,247 neurons detected within the larger subvolume. For the neurons, we extracted additional features from the somatic region of the cell, including its volume, surface area, and density of synapses. Dimensionality reduction on this feature space revealed a clear separation between neurons with well segmented somatic regions (n=69,957) from those with fragmented segmentations or sizable merges with other objects (n=12,290). Combining those features with the nucleus features, we trained a multi-layer perceptron classifier to distinguish excitatory from inhibitory neurons amongst the well-segmented subset, using the 80% of the manual labelled data as a training set, and 20% as a validation set to choose hyper-parameters. After running the classifier across the entire dataset, we then tested the performance by sampling an additional 350 cells (250 excitatory and 100 inhibitory). We estimate from this test that the classifier had an overall accuracy of 97% with an estimated 96% precision and 94% recall for inhibitory calls.

## Acknowledgments

The authors thank David Markowitz, the IARPA MICrONS Program Manager, who coordinated this work during all three phases of the MICrONS program. We thank IARPA program managers Jacob Vogelstein and David Markowitz for co-developing the MICrONS program. We thank Jennifer Wang, IARPA SETA for her assistance.

The work was supported by the Intelligence Advanced Research Projects Activity (IARPA) via Department of Interior/ Interior Business Center (DoI/IBC) contract numbers D16PC00003, D16PC00004, D16PC0005, and 2017-17032700004. The U.S. Government is authorized to reproduce and distribute reprints for Governmental purposes notwithstanding any copyright annotation thereon. HSS also acknowledges support from NIH/NINDS U19 NS104648, NIH/NEI R01 EY027036, NIH/NIMH U01 MH114824, NIH/NIMH U01 MH117072 NIH/NINDS R01 NS104926, NIH/NIMH RF1 MH117815, NIH/NIMH RF1 MH123400, and the Mathers Foundation, as well as assistance from Google, Amazon, and Intel. XP acknowledges support from NSF CAREER grant IOS-1552868. XP and AT acknowledge support from NSF NeuroNex grant 1707400. AT also acknowledges support from National Institute of Mental Health and National Institute of Neurological Disorders And Stroke under Award Number U19MH114830. RCR acknowledges support from NSF NeuroNex 2 award 2014862.

We thank John Philips, Sill Coulter and the Program Management team at the AIBS for their guidance for project strategy and operations. We thank Hongkui Zeng, Ed Lein, Christof Koch and Allan Jones for their support and leadership. We thank the Manufacturing and Processing Engineering team at the AIBS for their help in implementing the EM imaging and sectioning pipeline. We thank Brian Youngstrom, Stuart Kendrick and the Allen Institute IT team for support with infrastructure, data management and data transfer. We thank the Facilities, Finance, and Legal teams at the AIBS for their support on the MICrONS contract.

We thank Stephan Saalfeld, Khaled Khairy and Eric Trautman for help with the parameters for 2D stitching and rough alignment of the dataset. We thank Zane Hanson and Justin Singh for their contribution to manual matching of functional units to EM nuclei, and Donnie Kim for his contribution to pupil tracking. We also thank Rajkumar Raju for his contribution to parametric stimuli development. We thank Aaron Mok and Dimitri Ouzounov for their contribution to three-photon imaging development.

We thank Garrett McGrath for computer system administration, and May Husseini and Larry and Janet Jackel for project administration at Princeton University.

We thank Sandy Hider, Tim Gion, Derek Pryor, Dean Kleissas, Luis Rodriguez, Miller Wilt and the team from the John Hopkins University Applied Physics Laboratory (APL), as well as Marysol Encarnación and Martha Cervantes from the CIRCUIT Program at APL for supporting data assessments on the neural circuit reconstruction and data infrastructure through the Brain Observatory Storage Service & Database (BossDB, https://bossdb.org/, NIH/NIMH R24 MH114785).

We thank Frances Chance, Brad Aimone, and everyone at Sandia National Laboratories for their support and assistance.

We also would like to thank the following individuals for their work proofreading neurons in the MICrONS dataset: Natalie Smith (24101 edits), Divya Panchal (19384 edits), Michael Cook (17088 edits), Chris Ordish (14333 edits), Niyati (13897), Zainab Sorangwala (13777 edits), Nirali (13317 edits), Sholka (11569 edits), Krupali Shah (10570 edits), Dhwani Patel (10368 edits), Erika Neace (10312 edits), Dhara (9871 edits), Anuja (9337 edits), Zeba (8742 edits), Anjali Rajput (8674 edits), Claire Smith (8281 edits), Hemal (8084 edits), Harshil (8022 edits), Christopher Knecht (7199), Swagat Pal (7036 edits), Dhruvi Rami (6850 edits), Sweksha (6766 edits), Priyanka (6485 edits), Yashvi (6306 edits), Frank (5711 edits), Kavya Raval (5638 edits), Dhairya Dalal (5597 edits), Emily Phillips (5454 edits), Hetvi (5358 edits), Yuvaraju (4787 edits), Gary Hopkins (4505 edits), Neha (2968 edits), Anjali Pandey (2426 edits), Vaishakhi (2227 edits), Twinkal (1270 edits), Dylan Parodi (1007 edits), Kinjal (980 edits), Rachel Xu (947 edits), Kashish (911 edits), Clara Moore (828 edits), Vivia Lung (804 edits), Sarah Wu (746 edits), Taylor Gaito (643 edits), Jesse Gayk (570 edits), Lydia Fozo (506 edits), Diksha (438 edits), Mahaly Baptiste (416 edits), Ellie Macgregor (383 edits), Elanine Miranda (354 edits), Shruthi Bare (220 edits).

We would like to thank the “Connectomics at Google” team for developing Neuroglancer and computational resource donations, in particular J. Maitin-Shepard for authoring neuroglancer and help creating the reformatted sharded multi-resolution meshes and imagery files used to display the data. We would like to thank Amazon, the AWS Open Data Program, and the AWS Open Science platform for providing data and computational resources. We’d like to also thank Intel for their assistance.

We thank the Allen Institute for Brain Science founder, Paul G. Allen, for his vision, encouragement and support. Disclaimer: The views and conclusions contained herein are those of the authors and should not be interpreted as necessarily representing the official policies or endorsements, either expressed or implied, of IARPA, DoI/IBC, or the U.S. Government.

## Author Contributions

Conceptualization: HSS, FC, RCR, NMdC, AST, JR, XP

Methodology: JAB, MAC, SD, AH, ZJ, CJ, NK, KK, KLe, KLi, RL, TM, EMit, SMu, SSM, BN, OO, SP, WS, NLT, RW, WW, JW, RY, CMSM, AB, FC, DB, JB, MT, RT, GM, DB, WY, RCR, NMdC, LE, DK, SK, TF, JR, PGF, SP, EF, CX, TW, FHS, DY EYW,, BC

Software: JAB, MAC, SD, AH, ZJ, CJ, NK, KK, KLe, KLi, RL, TM, EMit, SMu, SSM, BN, OO, SP, WS, NLT, RW, WW, JW, RY, CMSM, FC, DB, RT, GM, WY, LE, DK, TF, SP, EC, TM, CAB, JJ, LMK, VR, DX, JM

Validation: CAB, WGR, PKR, JM

Investigation: JAB, MAC, SD, AH, ZJ, CJ, NK, KLe, KLi, RL, TM, EMit, SMu, SSM, BN, OO, SP, HSS, WS, NLT, RW, WW, JW, RY, CMSM, AB, FC, DB, JB, MT, RT, GM, DB, WY, RCR, NMdC, LE, DK, TF, AST, JR, PGF, SP, EF, SP, EC, TM, XP

Resources: ZHT

Data curation: SD, JG, JH, SK, MM, BS, SW, KW, SY, CMSM, AB, FC, JB, MT, NMdC, CG, GW, PGF, SP, EF, EC, SMc, MB, EMir, FY, AW, EJ, CZ, CAB, JJ, LMK, PKR, VR, BW, DX

Writing - Writing - Original draft: SD, TM, HSS, CMSM, FC, RCR, NMdC, AST, JR, PGF, SP, CP, BC, XP

Writing - Review & editing: JAB, MAC, AH, ZJ, CJ, NK, KLe, KLi, RL, EMit, SMu, SSM, BN, OO, SP, WS, NLT, RW, WW, JW, RY, AB, DB, JB, MT, RT, GM, DB, WY, LE, DK, TF, EF, SP, CX, TW, EC, CLS, AR, TM, PKR, JJ, DX, CAB, BW

Visualization: SD, AS, FC, NMdC,

Supervision: HSS, SY, FC, RCR, NMdC, AST, JR, XP, WGR, LMK, PKR, BW, DX

Project administration: TM, NMdC, SS, JR, RY, WGR, BW

Funding acquisition: HSS, RCR, NMdC, AST, JR, XP, BW

## Supplemental Data

**Supplemental Figure 1.**
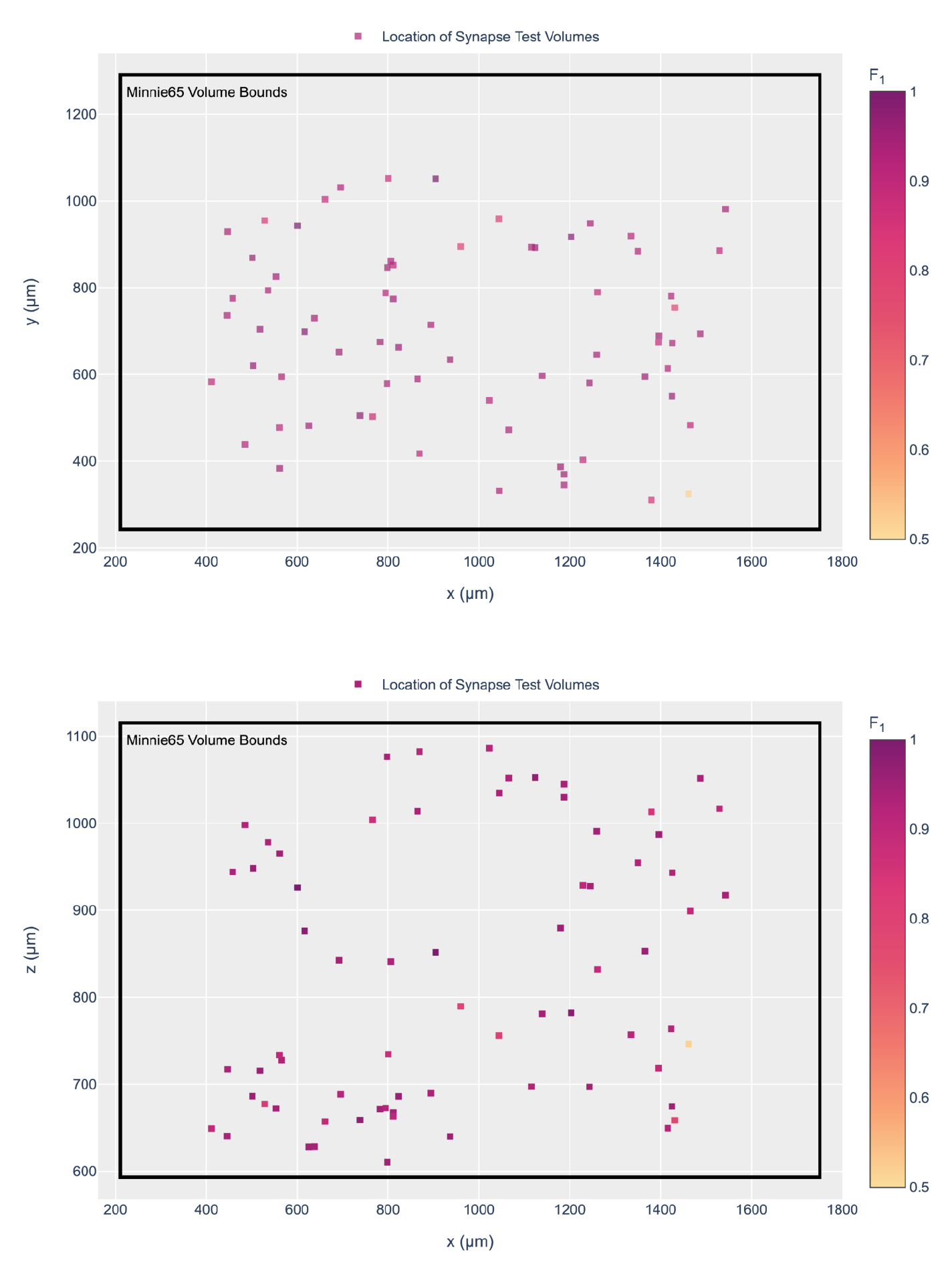
Test Volume Locations for Validation of Automated Synapse Detection. Location and distribution of test subvolumes (x=5.5µm, y=5.5µm, z=5.5µm) throughout the whole subvolume 65 that were used for validation of automated synaptic contact segmentation. Identification and annotation of synaptic contacts (n=8,611 synapses) were performed manually within each subvolume and compared with automated results to calculate subvolume and combined precision (96%), recall (89%), and F1 scores (98%), with test subvolume F1 scores visualized by color within each plot. The two panels show a coronal (**a**) and top (**b**) view of the location of the sampling sites. In (**a**) the vertical axis represents the pia to white matter direction and the horizontal axis represents the medial-lateral direction. In (**b**) the vertical axis represents the anterior-posterior direction and the horizontal axis represents the medial-lateral direction.

**Supplemental Figure 2.**
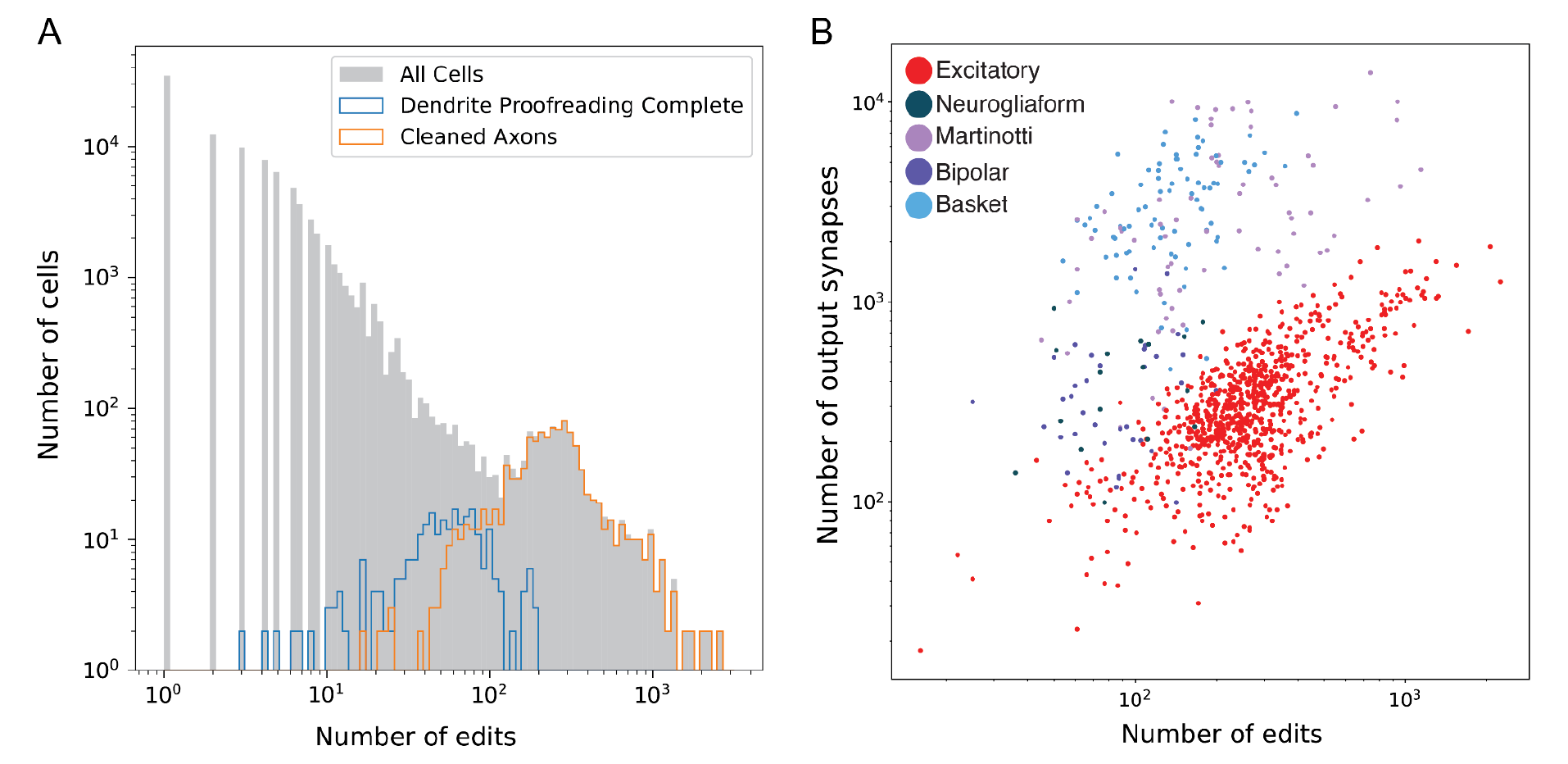
Proofreading statistics across the volume. **(A)** Histogram of number of edits across all the objects associated with nuclei. Distribution of neurons with complete dendritic proofreading highlighted in blue, and neurons with clean axons in orange. Cells with complete dendritic proofreading have often had some axon edits as well, so this is an upper bound on the number of edits required to fully extend dendrites. Most cells have had very little proofreading and have been mostly touched by automated methods. Note plot is on a log-log scale. **(B)** Number of edits compared to number of output synapses in reconstruction. For all the clean axons for which we have cell type annotations, the number of edits versus the number of outputs is plotted on a log log scale. Data points are colored with respect to their broad cell class. Generally, more extensively reconstructed axons have more edits, but there are also strong cell and cell-type specific effects. This reflects systematic differences in the thickness of axons of different cell types, as well as variation in how much of the axon is contained within the volume and the quality of the segmentation in different locations in the dataset.

**Supplemental Figure 3.**
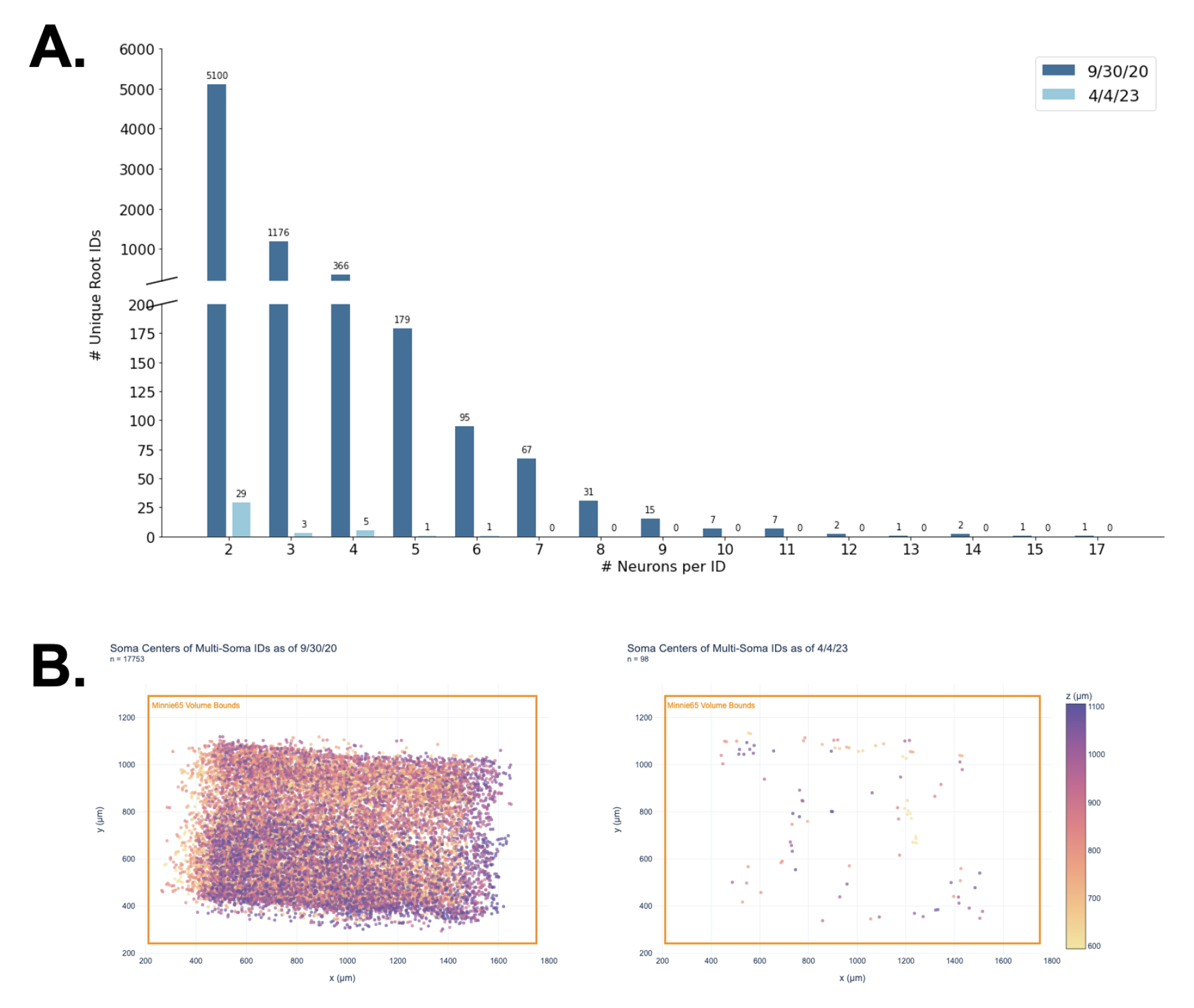
Multi-Soma Proofreading by APL. (A) Distribution of multi-soma IDs by number of neuronal nuclei was monitored throughout proofreading. Difference in multi-neuron root IDs before APL proofreading (dark blue) and after (light blue). Note that this shows the number of neurons per ID, which means that non-neuron somas are not counted. This figure was derived using the soma classification table: nucleus_ref_neuron_svm. Note that a small number of multi-soma IDs were skipped during APL proofreading because they contain low quality neurons merged to myelinated axons or they were falsely classified as neuronal (e.g. blood vessels); (B) Spatial distribution of multi-neuron ID soma centers (soma locations of merged cells containing ≥ 2 neuronal nuclei) before APL proofreading and after. Both are a lateral view of the volume that shows distribution across layers, from pia (top) to white matter (bottom). Color-bar represents depth.

**Supplemental Figure 4.**
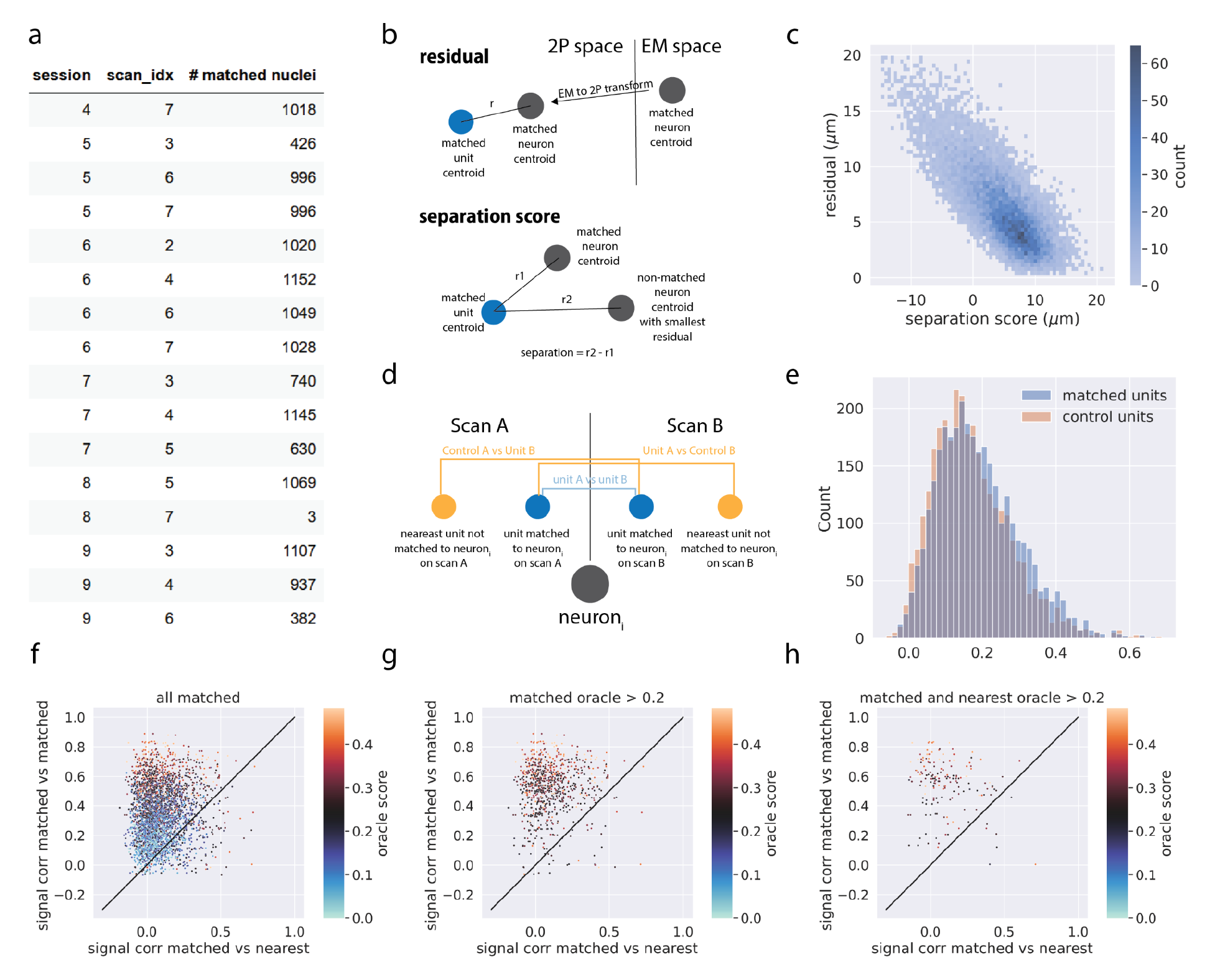
Manual Co-registration Metrics. **a)** the number of matched neuronal nuclei by session/ scan **b)** Schematic of the residual and separation score metric. **Residual:** For a matched EM nucleus to a functional unit, the residual is computed as the euclidean distance between the nucleus centroid and unit centroid after transforming the nucleus centroid from EM to 2P space with the spline-based coregistration. **Separation score**: For a matched EM nucleus to a functional unit the separation score is computed as the difference between the residual of the matched pair and the residual of the nearest EM neuronal nucleus that was not matched to the unit. Negative separation indicates that the nearest nucleus to the functional unit after coregistration was not chosen by the matchers. **c)** 2D histogram of separation score and residual. **d)** Schematic of in vivo signal correlation analysis. For every nucleus that was matched to at least 2 scans, and for every pair of scans, the in vivo signal correlation (correlation between trial-averaged responses to oracle stimuli) was computed between the matched unit in scan A to the matched unit in scan B. In addition, for each nucleus, two control correlations were computed, the matched unit in scan A to the nearest unit not matched to the same nucleus in scan B, and vice versa. **e)** The distribution of oracle scores for matched units and the nearest unit controls.**f)** scatter plot of signal correlations for all matched units (y-axis) vs the signal correlations for the nearest unit controls (x-axis) and colored by oracle score. Note that each matched unit pair has two data points on the plot for each of the two control correlations. **g)** same data as in **f)** restricted to matched units with oracle >0.2. **h)** same data as in **f)** restricted to matched units and control units with oracle > 0.2

**Supplemental Figure 5.**
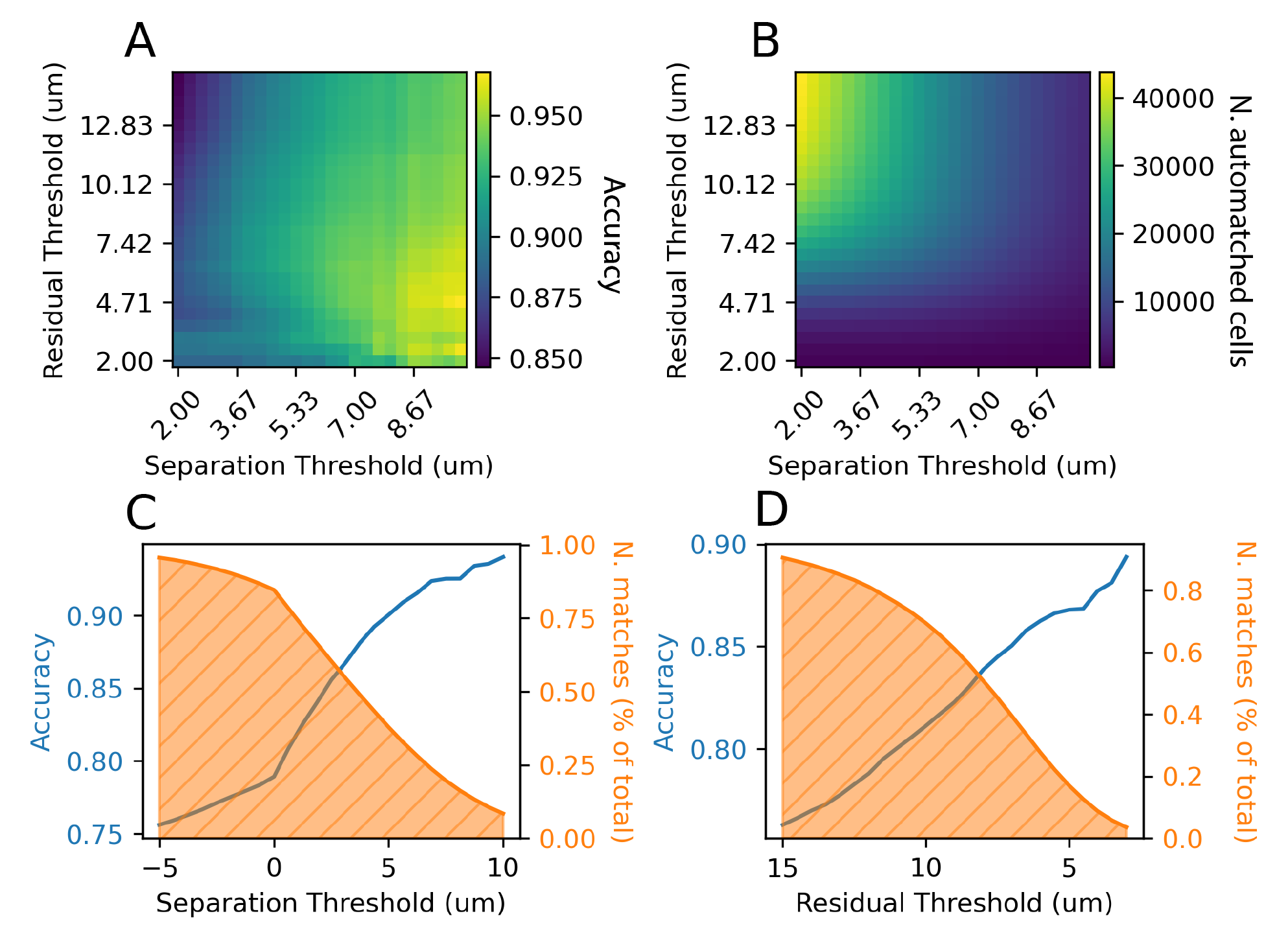
Fine-tuned coregistration. **a)** Agreement with manual matchers as a function of the residual and separation of the match after linear sum assignment. Note: separation does not depend on the output of the sum assignment and is calculated purely from Euclidean distances between centroids in each coordinate space. **b)** Number of matches that remain valid after imposing the threshold corresponding to the separation and residual thresholds. **c)** Accuracy and number of matches remaining as the separation threshold increases. **d)** Accuracy and number of matches remaining as the residual threshold decreases.

**Supplemental Table 1.** **Data Access Tools.** A list of programmatic open source tools for data access, what data products they help you access, and a link to their open source repository where there are further instructions and examples.

| Tool Name | Data Products | URLs |
| --- | --- | --- |
| cloud-volume | EM imagery, voxel segmentation and meshes | https://www.github.com/seung-lab/cloud-volume |
| MeshParty | EM meshes and skeletons | https://github.com/sdorkenw/MeshParty |
| CAVEclient | EM structural annotations (synapses, cell-types, etc), segmentation edits, local geometric properties | https://github.com/seung-lab/CAVEclient |
| microns-nda | Functional imaging DataJoint database | https://github.com/cajal/microns-nda-access |

**Supplemental Table 2.**
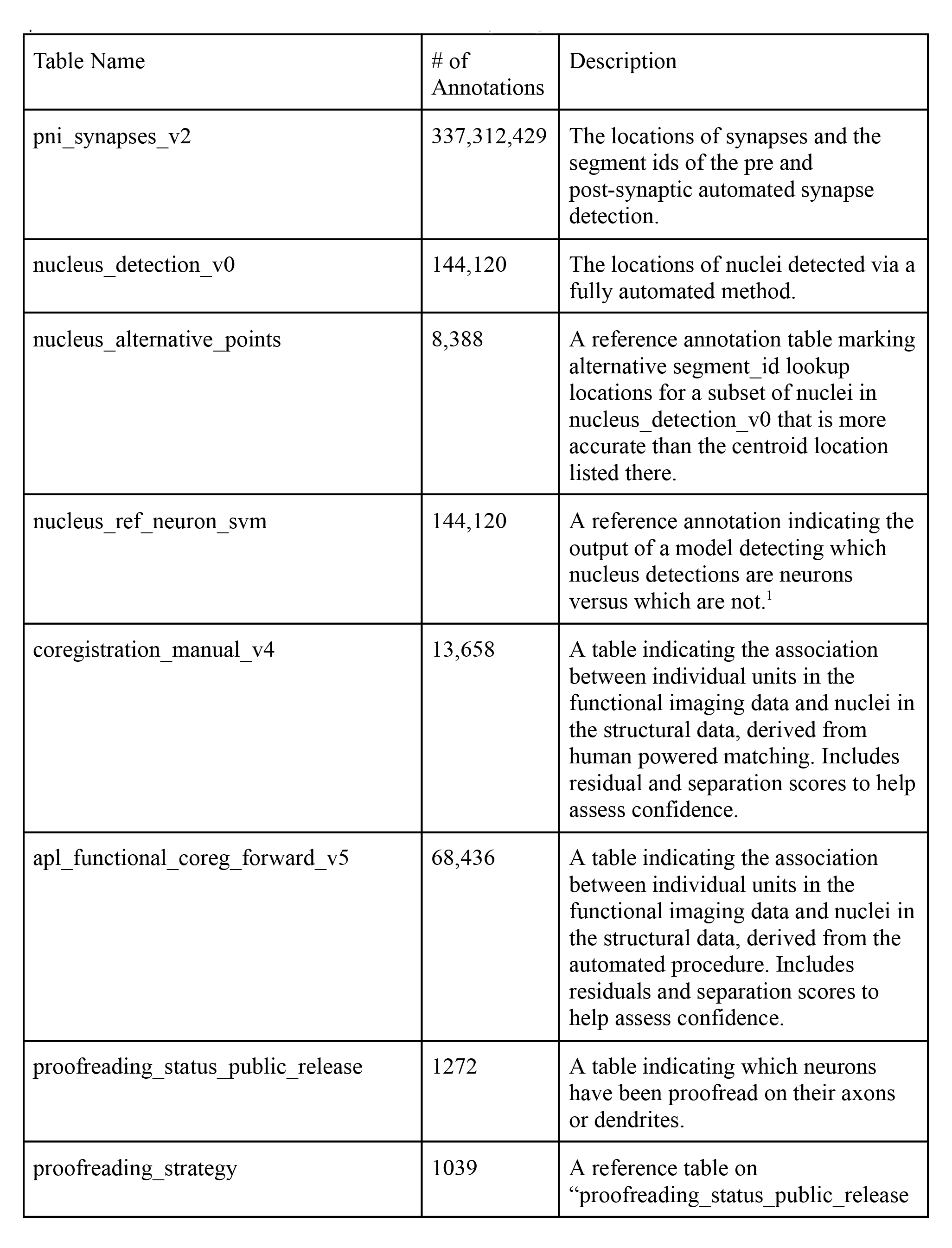
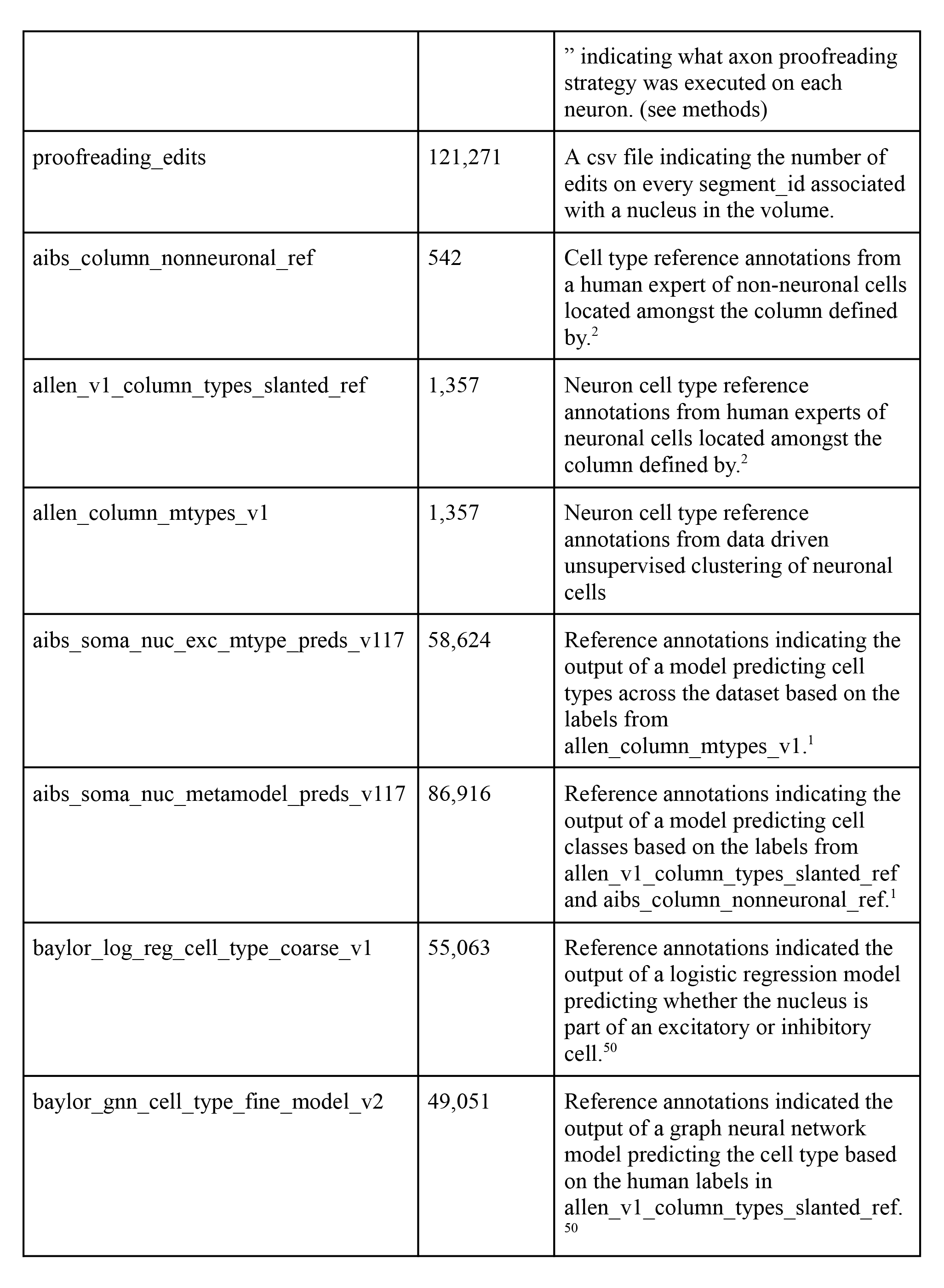
**CAVE Tables.** List of annotation tables that are part of the public release. Each table can be queried via the CAVE client and downloaded as a CSV from publicly available locations.

